# Rapid Assessment of T-Cell Receptor Specificity of the Immune Repertoire

**DOI:** 10.1101/2020.04.06.028415

**Authors:** Xingcheng Lin, Jason T. George, Nicholas P. Schafer, Kevin Ng Chau, Michael E. Birnbaum, Cecilia Clementi, José N. Onuchic, Herbert Levine

## Abstract

Accurate assessment of TCR-antigen specificity at the whole immune repertoire level lies at the heart of improved cancer immunotherapy, but predictive models capable of high-throughput assessment of TCR-peptide pairs are lacking. Recent advances in deep sequencing and crystallography have enriched the data available for studying TCR-p-MHC systems. Here, we introduce a pairwise energy model, RACER, for rapid assessment of TCR-peptide affinity at the immune repertoire level. RACER applies supervised machine learning to efficiently and accurately resolve strong TCR-peptide binding pairs from weak ones. The trained parameters further enable a physical interpretation of interacting patterns encoded in each specific TCR-p-MHC system. When applied to simulate thymic selection of an MHC-restricted T-cell repertoire, RACER accurately estimates recognition rates for tumor-associated neoantigens and foreign peptides, thus demonstrating its utility in helping address the large computational challenge of reliably identifying the properties of tumor antigen-specific T-cells at the level of an individual patient’s immune repertoire.

## 1 Introduction

The advent of new strategies that unleash the host immune system to battle malignant cells represents one of the largest paradigm shifts in treating cancer and has ushered in a new frontier of cancer immunotherapy [1]. Various treatments have emerged, including checkpoint blockade therapy [2, 3, 4], tumor antigen vaccine development [5, 6], and the infusion of a donor-derived admixtures of immune cells [7]. A majority of successful treatments to-date rely on the anti-tumor potential of the CD8+ T-cell repertoire, a collection of immune cells capable of differentiating between malignant cells and normal tissue by recognizing tumor-associated neoantigens (TANs) detectable on the cell surface [8]. Therefore, accurately assessing a T-cell repertoire’s ability to identify cancer cells by recognizing their tumor antigens lies at the heart of optimizing cancer immunotherapy.

A complete understanding of adaptive immune recognition and the tumor-immune interaction has remained a formidable task, owing in part to the daunting complexity of the system. For example, antigens and self-peptides contained in an epitope (i.e. recognizable peptide sequences) space of size ~ 20^9^ are presented to ~ 10^7^ unique T-cell clones in each individual [9], a small fraction of the upper limit of TCR diversity (~ 10^20^) [10, 11]. Moreover, their behavior is tempered via an elaborate thymic negative selection process in order to avoid auto-recognition [12, 13]. Here, T-cell clones, each with uniquely generated T-cell receptors (TCRs), interface with numerous (~ 10^4^) self-peptides presented on the major histocompatibility complex (p-MHC) of thymic medullary epithelial cells via TCR CDR3*α* and *β* chains, and survive only if they do not bind too strongly [14, 15, 16]. This process, together with systems-level peripheral tolerance [17, 18], imparts T-cells with durable tolerance to major self-peptides and influences many of the recognition properties of the resultant repertoire. The complexity of the adaptive immune system has attracted numerous mathematical modeling efforts quantifying the mechanisms underlying T-cell immune response. Collectively, the field has made significant progress in understanding the population-level effects of tolerance on T-cell recognition and self vs. non-self discrimination [14, 19]. This includes the T-cell repertoire’s effectiveness at discerning tumor from self-antigens [20], its ability to impart immunity against current and future threats [21, 22], and the extent of selection pressure that it exerts on an evolving cancer population [23, 24].

Any attempt at better understanding these system-scale properties must start with a reliable method to evaluate the interaction between specific TCR-p-MHC pairs. Despite this, a comprehensive, biophysical model capable of learning the energy contributions of each contact pair in a TCR-p-MHC system and applying them to new predictions remains elusive. To-date, experimental research has integrated solved crystal structures [25, 26] with peptide sequencing [27, 28, 29] to probe the physiochemical hallmarks of epitope-specific TCRs. Publicly available crystal structures have enabled researchers to identify detailed structural features that influence the binding specificity of TCR-p-MHC pairs, and machine learning algorithms have made progress on the complementary task of accurately predicting peptide-MHC binding [30, 31, 32, 33, 34, 35, 36] as well as TCR-peptide binding [37, 38]. However, the limited number of available structures relative to the diversity in MHC alleles and TCR-peptide combinations complicates extrapolation to unsolved systems. Alternate template-based structural modeling [39] and docking [40] approaches are limited by calculation speeds (at best one structure per minute), thus it is unlikely in the foreseeable future that such strategies can be used to investigate the number of TCR-peptide interactions necessary to study the problem at the immune-repertoire level, as this task easily requires the assessment of more than 10^9^ pairs simultaneously [16]. Prior attempts have approximated binding affinity by implementing statistical scores calculated from docking algorithms [40]. These scores are trained using examples of generic protein binding and thus lose the unique aspects of the TCR-peptide interactions.

To deal with this challenge, we develop a systematic TCR-p-MHC prediction strategy for rapid and accurate assessment of TCR specificity. Our strategy, which we refer to as the Rapid Coarsegrained Epitope TCR (RACER) model, is capable of differentiating between self and foreign antigens and can evaluate 10^9^ TCR-peptide pairs in the setting of TCR-peptide combinations restricted to a single MHC allele. This method we develop employs supervised machine learning on known TCR-peptide structures and experimental data to derive a coarse-grained, chemically-accurate energy model governing TCR-p-MHC interaction. This strategy was adapted from earlier efforts to predict protein folding [41, 42, 43, 44, 45, 46] and to screen the binding of small molecules [47, 48]. The MHC loci, while polymorphic, bind comparable numbers of peptides across various alleles [49]. Our calculations are restricted to a fixed MHC allele, but could be generalized with the use of additional training data. Confining our predictions to TCRs with a given MHC restriction enables the transferability of the method to TCRs that are not included in the training set. The approach provides a tractable means to extract pertinent TCR-peptide interactions so that affinity may be predicted based on similarly restricted TCR-peptide primary sequence data. RACER accurately distinguishes binding peptides across various TCRs and validation tests. Lastly, as a preliminary test of the usefulness of our approach, we simulate thymic selection and show agreement with previously established estimates of T-cell binding energy distributions, tumor neoantigen and foreign peptide recognition rates for a given class of MHC-restricted TCRs [50, 51]. Our *in silico* results share several features observed in experimental data including the degree to which post-selection TCRs preferentially recognize foreign antigen and TANs, in addition to the sequence diversity of epitope-specific TCRs. [52, 28]. Taken together, our results demonstrate RACER’s utility in learning the interactions relevant for high-throughput TCR-epitope binding predictions.

## 2 Results

### 2.1 RACER can distinguish peptides that bind strongly to a given TCR from those that bind weakly

The RACER optimization protocol (Fig. 1A) utilizes high-throughput deep sequencing data on TCR-peptide interactions across a large peptide library [27], together with known physical contacts between TCRs and peptides obtained from deposited crystal structures [53]. The training data comes from cases where all the peptides are displayed by the same allele of the mouse MHC class II molecule. The binding energy between TCRs and peptides, calculated based on a solvent-averaged coarse-grained pairwise model [46], was used as the metric to assess the TCR-peptide binding affinity. The interaction parameters for this solvent-averaged energy model were reoptimized here specifically for recognizing strong TCR-peptide interactions. Adapting an approach previously implemented for studying folding of proteins [54, 45], the RACER optimization strategy trains a pairwise energy model which maximizes TCR-peptide binding specificity. The energy model was optimized by maximizing the Z-score defined to separate the affinities of experimentally determined strong-binding peptides, called “strong binders” hereafter, from computationally generated, randomized decoys^1^. The optimized residue type-dependent energy model can then be used to calculate the binding energies of an ensemble of new TCR-peptide systems. As will be shown below, we performed three different levels of test (Fig. 1B), and find the predicted binding energies can differentiate strongly binding peptides from weak ones, provided they are displayed by the same MHC allele as that of the training set. Crucially, accurate predictions can be made even without knowledge of the actual crystal structure, although the predictions are improved when this additional information is available.

**Figure 1:**
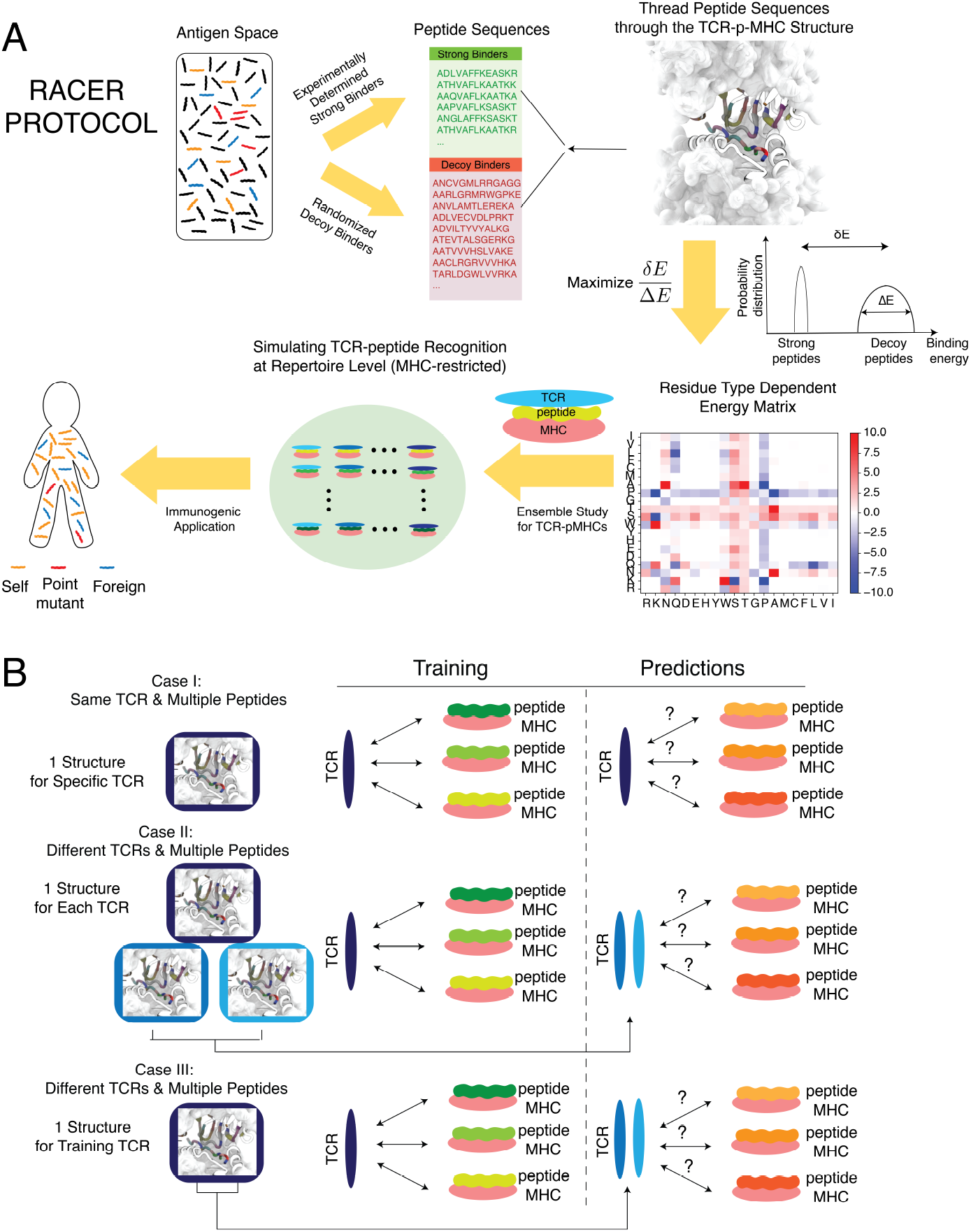
Summary of the modeling approach employed in this study. **A.** The optimization of RACER starts from a series of TCR binders obtained from the deep-sequencing experiments [27], as well as the corresponding TCR-p-MHC crystal structures deposited in the database [53]. The sequences of the strong binders, as well as the generated decoy binders from randomizing the non-anchoring sequences of the strong binders, are collected for parameterizing a pairwise energy model which maximizes the energetic gap between the strong binders and a randomized set of decoys. The resulting energy model can be used to quickly evaluate the binding affinities of an ensemble of TCR-peptide interactions at the population l evel. The calculated binding affinities can be used for simulating the negative selection process in the thymus, as well as measuring the recognition probability of the post-selection TCRs. Finally, this kind of ensemble study can be used for immunogenic applications that require input from an entire T-cell repertoire. **B.** Three tests were conducted to evaluate the performance of RACER. Case I: the training set includes one TCR-p-MHC structure and multiple peptide sequences. The test set includes the same TCR structure and a separate set of peptide sequences. Case II: the training set includes one TCR-p-MHC structure and multiple peptide sequences. The test set includes two different TCR structures (restricted on the same MHC allele) and two separate sets of peptide sequences. Structures for the two additional test TCRs are included in predictions. Case III: The training set includes one TCR-p-MHC structure and multiple peptide sequences. The test set includes only the sequences of two different TCRs (restricted on the same MHC allele) and two separate sets of peptides. Only the structure from the original training TCR was used in prediction (The interactions of interest are indicated by double-sided

Fig. 2 summarizes RACER’s predictive performance for a specific TCR (Case I in Fig. 1B). For this fixed TCR, pre-identified strong binding peptides and decoy peptides with randomized sequences were used to train the energy model (See Methods section for details). Another set of peptides independently verified experimentally as weak binders constitutes the testing set. The resulting energy model was then applied to calculate binding energies for the strong binders in the training set as well as the peptides in the testing set. This approach was repeated on three independent TCRs that are associated with the IE^k^ MHC-II allele: 2B4, 5CC7 and 226 (Details of these three TCRs are provided in Table S1). Although the experimentally identified weak binders were omitted from the training set, RACER effectively resolves binding energy differences between experimentally determined strong and weak binders having Z-scores, calculated in an analogous way as above by replacing decoys with experimentally-determined poor binders, larger than 3.5 in all cases (Fig. 2A), thus highlighting the predictive power of this approach.

**Figure 2:**
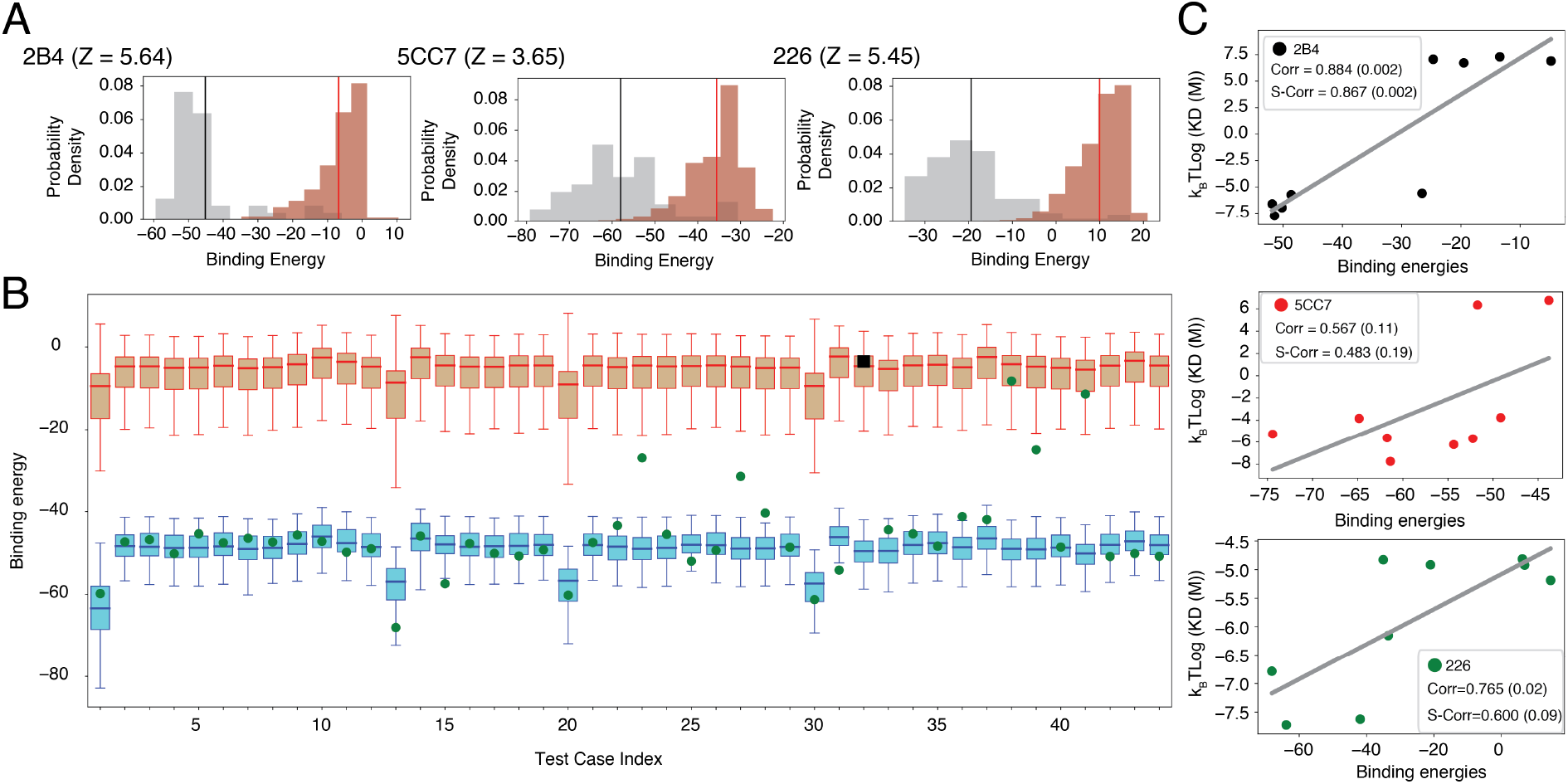
RACER can fully separate the strong binders of a specific TCR from its weak binders. **A.** For three TCRs (2B4, 5CC7 and 226) whose strong and weak binders have been experimentally determined [27], the RACER-derived calculated binding energies can well separate strong binders from weak ones of each indi-vidual TCR. **B.** In the leave-one-out-cross-validation exemplified using TCR 2B4, RACER can successfully recognize the withheld strong binders in 43 out of 44 tests, where the predicted binding energies of the with-held test binder (green) is lower than the median (red bar) of the experimentally determined weak binders. The only exception is marked as a black square. The whiskers are placed at the first and last datum points that fall within (m, M), where m = Q1 - 1.5IQR and M = Q3 + 1.5IQR, IQR = Q3 - Q1 represents the interquartile range. **C.** In a completely independent testing data measured by surface plasmon resonance (SPR) [27], the calculated binding energies of testing peptides correlate well with their experimentally determined dissocia-tion constant K_d_. Best-fit linear regression is depicted for each case. Corr: Pearson correlation coefficient. S-Corr: Spearman’s rank correlation coefficient. The p-value of each correlation coefficient is reported in the parenthesis.

Despite their relative sparsity in antigen space, strong binders play a central role in T-cell epitope recognition. It is obviously more difficult to predict strong binders than weak binders. To test RACER’s ability to identify strong binders, we performed a leave-one-out cross-validation (LOOCV) test, using data from TCR 2B4 as an example. For each test iteration, one known strong binder was withheld from the training set of 44 strong binders. Our optimization protocol was applied to train the energy model by using the remaining 43 peptides and then predicting the binding energy of the withheld peptide. This prediction was then compared to predicted binding energies of known weak binders, and the procedure was repeated for each of the 44 peptides. Our model is able to accurately distinguish the withheld strong binder in 43 cases (Fig. 2B). This is in stark contrast to a cluster-based attempt at strong binder identification based on peptide sequences alone, which at best correctly identifies 19 out of 44 strong binders (Supplementary note S1). The same LOOCV test was performed for TCR 5cc7 and 226, which correctly identified 120 out of 126 strong binders of 5cc7, and 267 out of 274 strong binders of 226. To further test the limit of RACER in detecting strong binders that have a more diverse sequence coverage, we performed a more demanding set of hold-out tests on a more comprehensive set of data from [27]. RACER can recognize peptides sharing little to no sequence identity with the native peptide (Figs. S1, S3), and is still able to recognize strong binders when a substantial portion of the training data is withheld (Supplementary note S2, S3 and Fig. S2, S3).

In order to further characterize RACER’s predictive power, an independent set of *K*_d_ values measured by surface plasmon resonance (SPR) [27] were compared with predicted affinities. The SPR experiments were performed on 9 independent peptide tests for each of the aforementioned three TCRs. RACER was used to predict the binding energies of each of those TCR-peptide pairs, each modeled with the structure of the corresponding TCR as the template. The free energies, *k*_B_*T* log(*K*_d_), were compared with calculated binding energies from RACER as a quantitative test of binding affinity prediction accuracy. Lower binding energies indicate stronger binding affinity so that a positive correlation between the *k*_B_*T* log(*K*_d_) values and calculated binding energies implies a successful prediction. As shown in Fig. 2C, RACER’s prediction of binding affinities for these 9 peptides correlates well with experimental measurement, with an average Pearson correlation coefficient of 0.74. The predicted order of binding affinities is also consistent with those from the experiment, with an average Spearman’s rank correlation coefficient of 0.65.

### 2.2 RACER’s residue type-dependent interactions are optimized specifically for TCR-peptide recognition

The data utilized by RACER includes strong binders and an input crystal structure, as well as TCR and peptide primary sequences, which determine an interaction pattern that was then used to construct a system-specific force field. To illustrate this, we focus on the 2B4 TCR as an example (Fig. 3). The crystal structure of TCR 2B4 (Fig. 3A) reveals that there can be many threonine (T) and asparagine (N) residues on the CDR loops region of the TCR. In the strong binder set, these residues tend to interact with specific peptide residues such as alanine (A), as seen for the specific peptide given in the figure. This notion can be formalized by showing the matrix of observed probabilities of close proximity of specific residue pairs. Thus, we see that certain pairs such as A-T and A-N are significantly enriched in the set of strong binders, while much less so in the decoy set (Fig. 3B). This then will mean that the optimized energy model shows the strongest attractions between the A-T, A-N residue pairs (Fig. 3C). This relative enrichment contrasts with the TCR tryptophan (W) residue which frequently interacts with alanine (A) in both strong binders and decoy peptides. As a result, the optimized energy model does not favor the A-W interaction.

**Figure 3:**
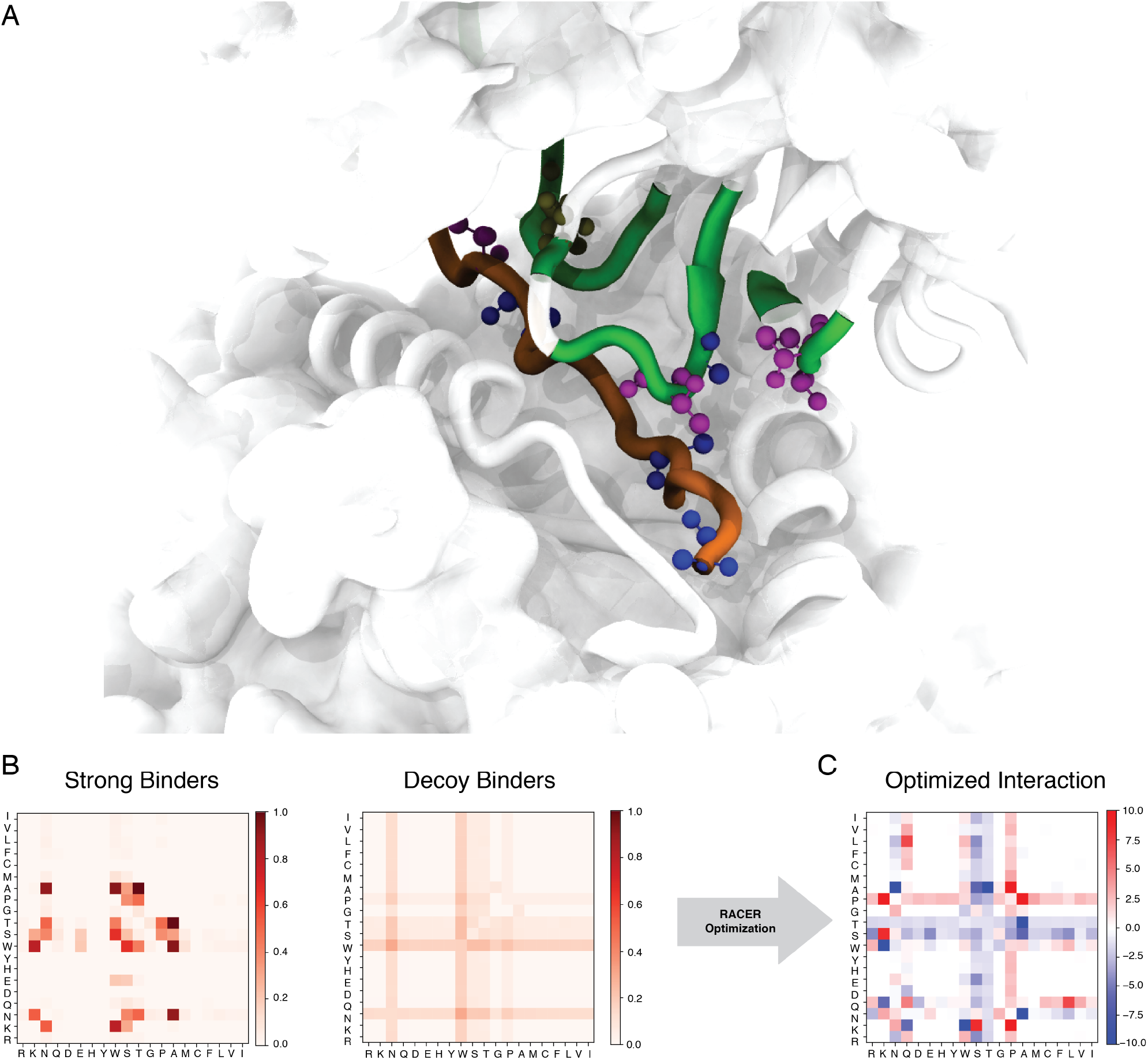
The specific contact pattern from the TCR-peptide structures dictates a optimized energy model different from those of a typical protein-folding force field. **A.** The 3D crystal structure of the 2B4 TCR bound to a specific peptide (PDBID: 3QIB). The parts of the structure that are in contact between the TCR and peptide are color-highlighted as green (TCR) and orange (peptide). Also shown are residues alanine (blue), threonine (magenta) and asparagine (tan) which are prevalent in this structure (CPK representation [60]). **B.** The probability of contact formation between each two of the 20 amino acids in the set of strong binders (left) and the set of randomized decoy binders (right) of TCR 2B4. **C.** The residue-based interaction strength determined by RACER for TCR 2B4. A more negative value indicates a stronger attractive interaction between the corresponding two residues.

This energy matrix is rather distinct from those typically used for studying protein folding. In order to compare the RACER-derived interaction matrix to well-established force fields described in the protein folding literature, we substitute for our interaction matrix either the standard AWSEM [46] (optimized on deposited folded proteins) force field or the Miyazawa-Jernigan (MJ) statistical potential [55] (constructed using the probability distribution of contacting residues from deposited proteins) and calculate the corresponding binding energy predictions for the TCR 2B4 peptides. We find that neither of them effectively resolves these groups, with Z-scores of 0.69 and 1.28, respectively (Supplementary note 4 and Fig. S4). Similar trends were observed utilizing the peptides corresponding to the 5CC7 and 226 TCRs, demonstrating the necessity of RACER’s *de novo* identification of pertinent structural information for studying the TCR-peptide system.

### 2.3 RACER’s interactions enable accurate predictions across various TCRs restricted to a given MHC allele

Given RACER’s accuracy in resolving test peptides presented to the specific TCR used for training, we next explored the feasibility of extending predictions to additional TCR-peptide pairs albeit with the same MHC restriction. Toward this end, we assessed whether the physical contacts implicitly encoded in RACER’s optimized force field were conserved within IE^k^-restricted TCR-peptide pairs. The three IE^k^-restricted TCRs considered in our analysis all have been tested with peptides bound to the IE^k^ mouse MHC molecule. The available crystal structures have a significant degree of structural similarity at the TCR CDR3-peptide binding interface (see Fig. 5 of [27]). We further quantified the TCR CDR3-peptide contacts for each pair, constructing a contact map based on their crystal structures (see Methods section for full details). Our results suggest that, despite differences in TCR and peptide primary sequences, the set of strong binding TCR-peptide pairs share common structural features which should aid in imparting transferability of the trained interaction matrix for accurate extrapolation to unknown pairs (Fig. 4). We find however that these features are not preserved across different MHC class II genes (Fig. S5), again indicative of the importance of incorporating structural information.

**Figure 4:**
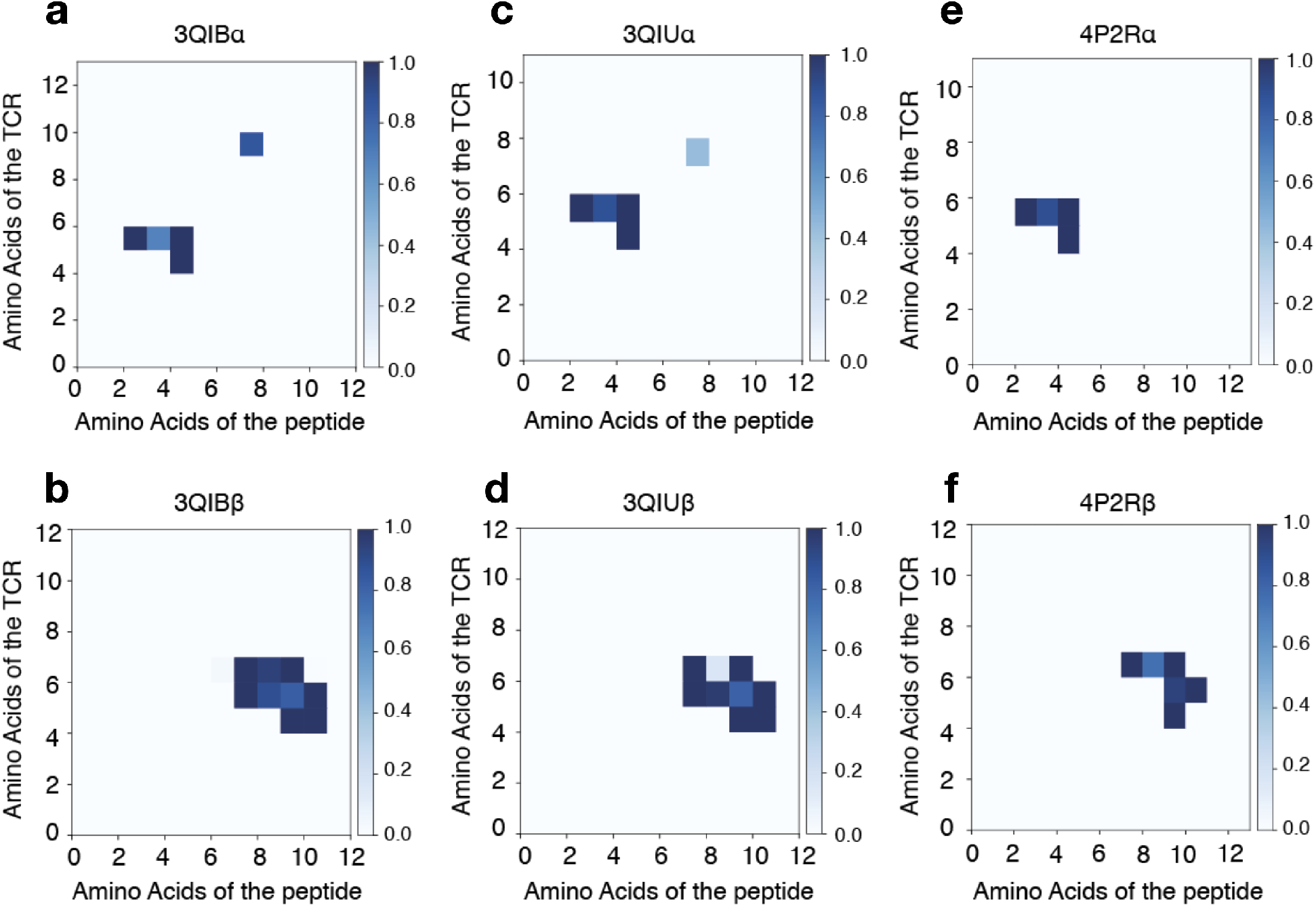
The contact maps of TCR-peptide pairs within the same MHCII allele share structural similarity. Contact maps are calculated using distances from each pairwise TCR-peptide amino acid combination using Eq. 6 for the following MHC-II IEk-restricted TCR-peptide pairs: 3QIB - peptide ADLIAYLKQATK with TCR 2B4 **A.** CDR3*α* (AALRATGGNNKLT) and **B.** CDR3*β* (ASSLNWSQDTQY) chains; 3QIU - peptide ADLIAYLKQATK with TCR 226 **C.** CDR3*α* (AAEPSSGQKLV) and **D.** CDR3*β* (ASSLNNANSDYT) chains; 4P2R - peptide ADGVAFFLTPFKA with TCR 5cc7 **E.** CDR3*α* (AAEASNTNKVV) and **F.** CDR3*β* (ASSLNNANSDYT) chains. Similarity in interaction topology across TCR-peptide pairs is observed by comparing the contact silhouette of interacting coordinates for the *α* (top row) and *β* (bottom row) TCR sequences.

**Figure 5:**
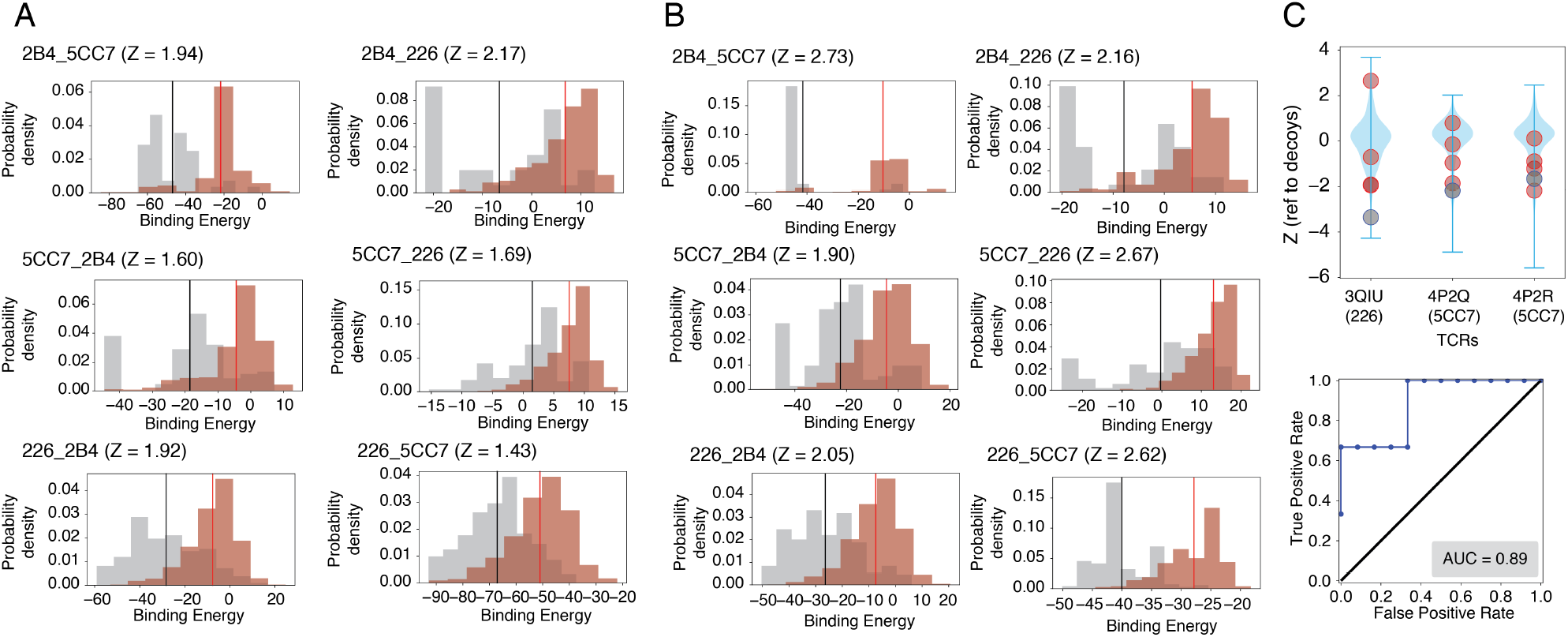
RACER shows transferability in terms of predicting TCR-p-MHC interactions across different TCRs. **A.** The energy model trained based on one TCR (e.g. 2B4) is capable of resolving the experimentally determined strong binders from weak binders of the other two TCRs (e.g., 5CC7 and 226). **B.** By adding strong binders from crystal structures of the other two TCRs into training sets, RACER can be further improved for identifying the experimentally determined strong binders. The title of each figures follows the format of “target training TCRs”, e.g., “2B4 5CC7” means using the energy model trained from TCR 5CC7 for predicting the peptide binding affinities of TCR 2B4. “Xtals” means the strong binders from the crystal structures of the other two TCRs were added into the training set. **C.** Upper panel: The energy model trained on TCR 2B4 is used to predict the binding energies of sequences from other TCRs associated with the IEk-associated TCRs [56]. Z-scores of known strong binders (grey) and weak binders (orange) provided by [56] were calculated referenced to a set of 1000 decoy peptides with randomized sequences (blue violin plot), with lower Z-scores indicating better predictive performance. Lower panel: The calculated Z-scores of each TCR were used to depict Corresponding ROC curve and AU-ROC (0.89, lower panel).

RACER’s ability to accurately identify strong binders based on training with a fixed TCR, together with the fact that a majority of the contact structure is preserved within a given MHC-restricted set of TCRs, suggested that we assess RACER’s ability to accurately predict binding peptides for other similarly restricted TCRs. Toward this end, we apply the energy model optimized using binding data for one of the three TCRs to predict the TCR-peptide binding energies of the remaining two holdout TCRs (Case II in Fig. 1B). To do this, we initially use a known structure for each of the holdouts, and the interaction matrix learned on the training TCR to predict the binding energies of the experimentally determined strong and weak binders for each of those holdout TCRs. Although the Z-scores measured for these alternate TCRs are lower than those found previously in Sec 2.1, RACER still successfully distinguishes a majority of strong binders from weak binders, with an average Z-score of 1.8 (Fig. 5A). This demonstrates that, despite CDR3 primary sequence diversity, distinct TCR-p-MHC systems associated with the same MHC allele still share similar structural-sequence patterns.

In order to test whether the incorporation of additional TCR structural information in the optimization step could improve RACER’s predictive accuracy, we next included crystal structures for the remaining TCRs (5cc7 and 226) together with a single strong binder for each case into the training set comprised of 2B4 peptide pairs (See Methods section for details). This procedure was repeated three times by substituting for the training set TCR and peptide pairs. We find that the new energy model demonstrates significant improvement in Z-scores. These results suggest that future incorporation of additional crystal structures of target TCRs may lead to improved resolution of strong and weak binders via refinement of the optimized energy model.

To provide an additional test and to quantify our discrimination capability, we used an independent dataset from [56]. Four independent TCRs (PDB ID: 3QIB, 3QIU, 4P2Q, 4P2R) from their curated benchmark dataset are associated with the IE^k^ allele; note that three of these overlap with the TCRs in our current study. To test the performance of RACER for different TCR-peptide pairs, we used the energy model trained based on 2B4 (3QIB) to predict the binding energies of both strong and weak binders for the three remaining TCRs. This calculation again uses the structure found for the one strong binding peptide for each of the 3 TCRs. Our calculation re-emphasizes that RACER can successfully distinguish strong binders even when it is trained based on a different TCR (Fig. 5C), with an AUC of 0.89. Of note, when we tested data from the same study involving TCR-p-MHCs with different MHC alleles, RACER cannot pick out strong binders, presumably due to the markedly different TCR-peptide interacting patterns (Fig. S5). As a more comprehensive test of RACER’s transferability, we included other TCR-peptide pairs from [56]. RACER is capable of recognizing the strong binders across TCRs with different V*α* and V*β* genes, and does so more effectively when there are multiple copies of TCR-peptide pairs available for training (Supplementary note S5, Fig. S6).

Next we address the question of the extent to which it is necessary to have at hand at least one TCR-p-MHC crystal structure in order to use RACER’s interaction matrix to identify other good binders (Case III in Fig. 1B). Of course to evaluate the binding energy we must have a structure; the alternative to having a measured structure for a new sequence is to thread that new CDR3 sequence into the crystal structure used for the training data, which potentially introduces an uncertainty in registration. For the cases at hand, this issue arises only for the *α* chain as the *β* chains for all three TCRs are all of length 12 and there is no residual ambiguity. We tested the simplest possible assumption, namely that we start at the same place where all three chains have the first two residues AA and leave no gaps. Fig. S8 shows that this procedure again leads to successful discrimination between good and poor binders, with an average Z-score of 2.36. As a comparison, the best performance of a recent sequence-based predictor trained by using artificial neural networks [38] can recognize the strong binders of TCR 5CC7, but not TCR 2B4 and 226 (Supplementary note S6 and Fig. S9). Similar tests were also performed for the TCR-peptide pairs from [56]. RACER still capably recognize the strong binders across TCRs with different V*α* and V*β* genes (Supplementary note S5, Fig. S7). Thus, we conclude that the structures are sufficiently similar that not only can we use the interaction matrix derived from a single TCR training set for other TCRs but we can also use the same structure. This then allows us to make estimates at the repertoire scale without the impossible task of creating extremely large numbers of TCR-p-MHC structures.

### 2.4 RACER-optimized T-cell repertoire binding assessment accurately represents thymic selection

Using RACER, we can determine general properties of TCR-p-MHC binding distributions and compare to empirical observations. These results highlight the advantage of a method capable of highthroughput analysis. The basic idea follows from the fact shown above that we can make reasonable assessments of binding strength by using only one structure and its associated interaction matrix. Our focus here is the process of negative selection and its effect on the surviving repertoire. Toward this end, we utilized the crystal structure of the 2B4 TCR-peptide contact region to create 10^5^ simulated TCRs and 10^4^ self-peptides by randomizing uniformly the CDR3 and peptide sequences over amino acid space. To avoid registration issues, we always choose simulated TCRs to have exactly the same number of *α* and *β* chain residues as does the 2B4 TCR. This was repeated using 10^4^ self-peptides and 2000 TCRs, this time weighting the CDR3-peptide interactions by each of the the three contact maps in Fig. 4. The same approach was applied to a model that assumes a strictly diagonal contact map motivated by previous analytical work [20], with randomization of the TCR sequence taken over all possible positions in the contact map.

A given TCR survives only if it binds to all self-peptides below a fixed activation threshold. The maximum binding energy over the set of self-peptides for each TCR defines a selection curve (Fig. 6A), which describes the percentage of negatively-selected T-cells as a function of the cutoff energy threshold. Selection curves for the three TCR sets using the contact maps in Fig. 4 utilized the RACER energy matrix and compare reasonably to the diagonal contact map motivated by previous analytical work (Fig. 6A red curve). While the variance in each case is similar, mean-shifts in each selection curve correlate directly with the number of contacts in the CDR3 *α* and *β* chains (Fig. 4). These findings further reinforce the relevance of TCR-p-MHC-specific structural interactions encoded in the RACER-derived energy potential for binding prediction and T-cell repertoire generation. Although empirical estimates of the percentage of surviving TCRs during thymic negative selection vary between 20% and 50% [57, 58, 59], we calculate relevant recognition behavior for all selection rates, restricting our analysis to 50%, when applicable.

**Figure 6:**
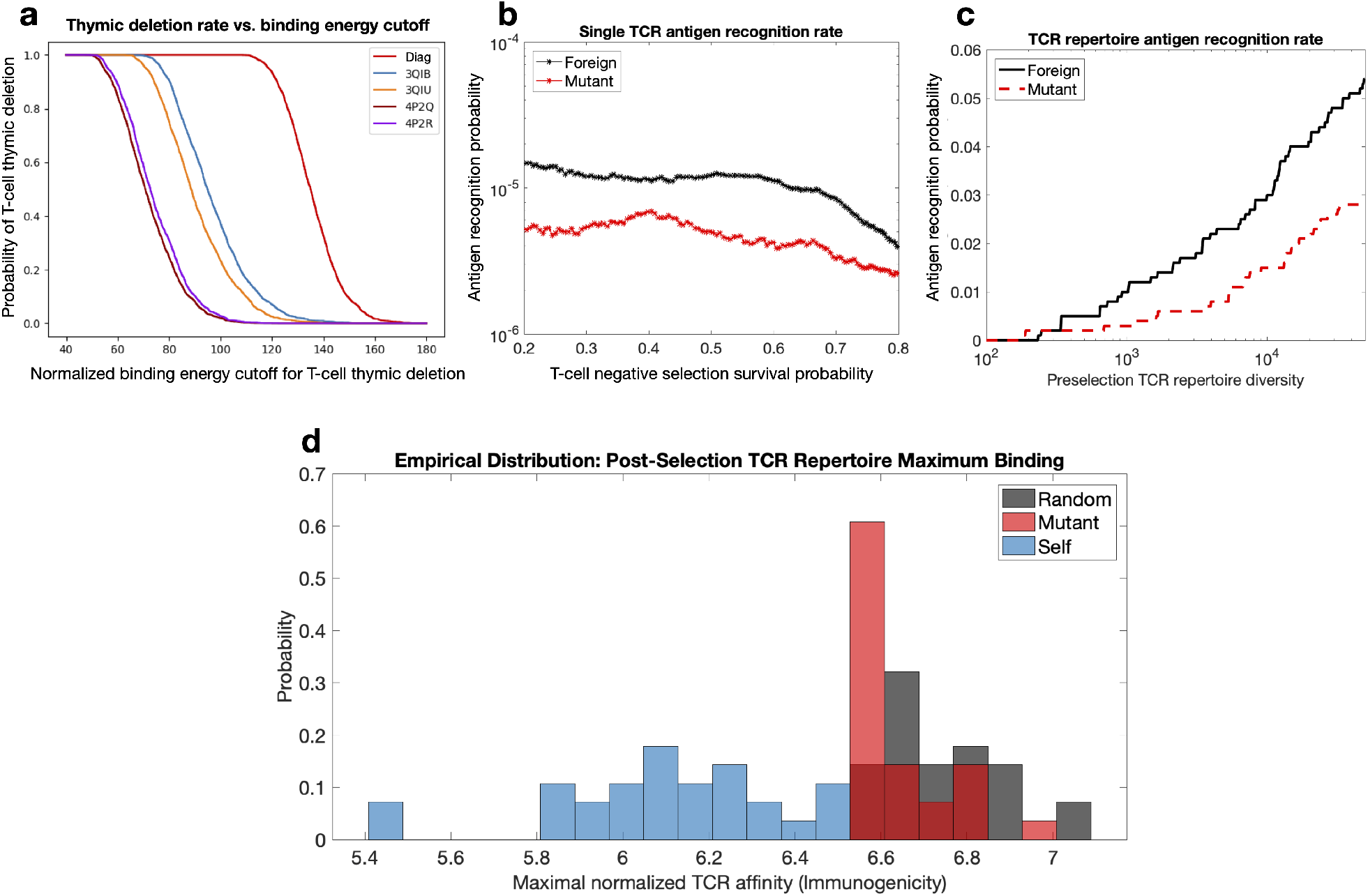
T-cell repertoire simulations of thymic selection and antigen recognition in the RACER model. RACER-derived simulations of TCR recognition exhibit sensible thymic selection and similarity in the recognition rates of foreign and point-mutated self antigens. **A.** Simulated thymic selection curves (T-cell recognition probability as a function of negative selection binding energy cutoff) incorporating the effects of non-adjacent contacts (given in Fig. 4) using *N_n_* = 10^4^ uniformly randomized self-peptides and *N_t_* = 2000 randomized IE*k*-restricted TCRs. 4P2Q and 4P2R (purple) use T-cells generated by randomizing the CDR3 region of TCR 5cc7, while 3QIB (blue) randomizes the CDR3 of TCR 2B4, and 3QIU (yellow) randomizes the CDR3 of TCR 226 (in all cases, randomized CDR3 lengths were unchanged from the original TCR) (red curve uses RACER energy using a diagonal contact map model whose study here is motivated by previous work [20]). **B.** Utilizing RACER-derived energy assessments from the 2B4 crystal structure, the probability of recognizing foreign and point-mutant antigens for individual post-selection T-cells is plotted as a function of the percentage of TCRs surviving negative selection (ordinate of the graph in panel a, simulations averaged over all post-selection TCRs with pairwise interactions amongst 10^3^ random peptides and 10^3^ point-mutant peptides). **C.** The recognition probability of foreign (black) and mutant (red) peptides by the entirety of the TCR repertoire is plotted as a function of pre-selection TCR repertoire diversity, with negative selection thresholds giving 50% survival. **D.** RACER-derived immunogenicity of foreign, mutant, and self antigen. The maximum binding affinity over all post-selection T-cells for immunogenic random (gray) and point-mutated self-peptides (red) is compared to that of thymic self-peptides (green) (There were 28 point-mutated peptides that had at least one T-cell recognition event. To keep an equal number of peptides in each distribution, we compared these with the top 28 similarly ordered foreign peptides and 28 randomly chosen self-peptide groups).

Most self-peptides present in thymic selection are expected to participate in the deletion of self-reactive T-cells. Previous work has suggested that this desideratum can be used to determine if a high-throughput model is behaving in a statistically sensible manner; specifically, a reasonable model of thymic selection would feature a majority of self-peptides contributing to the selection of immature T-cells. A rank order of these self-peptides based on their ability to recognize unique T-cells, or potency, characterizes the extent to which each self-peptide is utilized in thymic selection. The RACER-derived rank order using the 2B4-optimized data generates reasonable behavior with respect to this criterion (Fig. S10A).

One key issue influencing adaptive immune recognition of tumor-associated neoantigens (TANs) is the recognition efficiency of peptides closely related to self (e.g. point mutants) relative to foreign peptide recognition. The fact that the immune system can in fact be enlisted to attack tumors suggests that negative selection leaves intact the ability to bind strongly to tumor associated antigens. Comparison of a post-selection TCR’s individual recognition potential shows relatively minor differences between foreign and point-mutant self-peptides (Fig. 6B), with variances of these estimates overlapping with one another and in line with previous theoretical estimates (Fig. S10B). While individual recognition probability measure a single TCR’s ability to recognize antigen, repertoire recognition probability estimates a particular MHC-restricted post-selection repertoire’s ability to recognize antigen. An analogous comparison of the post-selection TCR repertoire recognition probability of foreign and mutant peptides demonstrates that this minimal difference is maintained at the aggregate immune system level (Fig. 6C). This then explains the observed ability of adaptive immune targeting of tumors in a manner that depends on the mutational load of the malignant cells.

Our prior theoretical model posited thymic selection as an optimization problem with a survival cutoff near 1*/e* resulting in the production of maximally efficient thymic selection [9, 20]. Calculating the product of survival and recognition probabilities yields a broad curve with large values located at intermediate survival cutoffs, including the previously predicted optimal survival cutoff (Fig. S10C). We also can compare our RACER-derived output to immunogenicity scores of experimentally-determined thymic self-peptides, foreign peptides, and TANs [52]. To accomplish this, we calculate the maximal binding energy of post-selection T-cells against each peptide class, a repertoire-level measure of immunogenicity. We find that our model is in broad agreement with other studies [52] that have placed the immunogenicity of TANs intermediary to that of foreign and self-peptides with a distribution closer to the foreign group (Fig. 6D).

For additional evaluation of RACER predictions against known experimental findings, we studied the similarity of T-cell CDR3 sequences at the repertoire level generated by RACER-derived TCR-p-MHC specificity maps. These were then compared to the properties of experimentally assessed TCR repertoires of known specificity against pre-identified antigens [28]. In our simulation, post-selection TCRs recognizing the top 10 foreign antigens were collected and clustered using a similar discrete Hamming metric that weighted the CDR3 sequences as in [28]. Dendrograms obtained from hierarchical clustering identified a diverse set of specific TCRs (Fig. S11A). Because our model only considered a small (10^5^) number of TCRs relative to the allowable diversity of CDR3 primary sequence space, we then augmented our T-cell repertoire by *in silico* site-directed mutagenesis to include 100 additional, closely related TCRs for each of the 10 aforementioned peptides, each undergoing identical selection and recognition tests. This augmented list of T-cells accurately recapitulates the empirical observation of a mixed set of specific T-cells, comprised of diverse and homologous clusters of TCR sequences (Fig. S11B), and demonstrates RACER’s utility for identifying largely diverse TCRs with specificity against a known antigen. Taken together, these results agree with previous studies and reinforce the utility of RACER for performing repertoire-level analyses.

## 3 Discussion

We have introduced RACER, an optimized molecular energy model that can be utilized to quickly assess TCR-peptide interactions and distinguish strong-binding pairs. RACER requires only ~0.02s for evaluating one TCR-peptide pair, thousands of times faster than available alternative approaches, while preserving reasonable prediction accuracy (Figs. 2, 5). Consequently, our method can be used to study large collections of MHC-restricted TCR-peptide pairs, enabling *in silico* studies of thymic selection and prediction of T-cell antigen recognition based on primary sequence data.

### 3.1 Specificity vs. Generality of the optimized energy model

The unique topology of the TCR-p-MHC structure encodes a system-specific residue-type dependent interaction matrix for TCR-peptide pairs. Significantly, the sequences and structures of TCR-peptide systems were found experimentally to be relatively conserved among various peptides [27, 28, 26]. The preserved sequence and structural features dramatically limit the physiochemical space explorable by TCR-peptide residue pairs. Moreover, since RACER is optimized on a TCR-peptide system, the arrangement of the contacts between TCR and its cognate peptide (Fig. 4) gives rise to a post-optimization energy model (Fig. 3) rather distinct from the traditional hydrophobic-hydrophilic interaction patterns [61] used for studying protein folding, such as the MJ potential [55]. This hypothesis is strongly supported by the observation that RACER is capable of identifying strong binders of corresponding TCRs (Fig. 2) while previous methods fall short (Fig. S4).

The departure of RACER from a typical protein-folding force field also results from the optimization performed for TCR-peptide systems. Because we are interested in resolving strong binders from weak ones with a finite dataset, our optimization distinguishes between these two sets of binders by enlarging their energetic gap in the training process. By maximizing the Z-score between strong and weak binders, RACER learns an effective binding energy which likely amplifies small difference in thermo-stability among candidate binders. Such amplification, however, affects neither the identification of the strong binders of a specific TCR nor the subsequent ensemble study of peptide recognition, since only the order of binding affinities among individual TCR-p-MHC pairs matters for our results.

### 3.2 Structural information from available crystal structures improves the predictive power of RACER

Our pairwise RACER model offers a novel avenue for developing models that incorporate information contained in available protein structures. Prior investigations have applied artificial neural networks for predicting strong binders of TCR [37, 38] and MHC [30] molecules based only on the primary sequences. Although deep learning can implicitly account for higher-order interactions, such approaches may still be limited by the available sequences that can be identified from experiments. RACER alleviates the high demands for primary sequences by including existing crystal structures in a pairwise potential. In order to provide a comprehensive characterization of RACER’s predictive power using each of our predictive assessments, our training set was limited to cases that had pre-identified TCR-peptide pairs given their known crystal structure [56]. While RACER effectively resolved strong and weak binders in all cases where the training and test peptide were identical, approximately half of test cases derived from this set contained training and test peptides that were dissimilar. In these cases alone, RACER correctly predicts 67% of the examples (Z-score > 1.0). The resulting predictive accuracy demonstrates that our structurally informed pairwise model is able to resolve TCR-p-MHC specificity in a majority of available test cases. Further experimental validation will be required to definitively assess RACER’s ability to resolve TCR-p-MHC specificity across all possible TCR-peptide pairs within a given MHC allele. This challenge remains a top priority for future investigations on repertoire-level TCR-peptide assessment.In designing RACER to achieve rapid and accurate predictions, our calculation only includes pairwise energetic interactions, while omitting contributions from conformational entropy. While RACER maintains reasonably high predictive accuracy, more accurate assessments of the TCR-p-MHC binding free energy will likely lead to improvements and is a focus of subsequent work.

In cases with available crystal structures, contact map analysis revealed a largely conserved interaction pattern reproduced across a variety of TCR-peptide pairs associated with the IE^k^ MHC II allele (Fig. 4), providing an explanation for the transferability of RACER-derived interactions when trained on a particular crystal structure. Moreover, these results contributed to variety in the selection behavior of individual TCRs in that TCR-peptide systems having more interactions in their corresponding contact map were correlated with systematic shifts in their mean binding energies, which subsequently correspond to differences in their post-thymic selection inclusion probability (Fig. 6). Previous investigations have characterized the probability distribution for generating particular TCR sequences in VDJ recombination, and have even suggested that the *a posteriori* observed post-selection TCRs had greater generation probabilities [15, 62], with so-called “public” TCR sequences being observed in multiple individuals. Incorporation of contact maps into our generative model contributes to variations in T-cell survival probability, and may offer a physical interpretation of why public repertoires may survive thymic selection at higher rates[63], in addition to providing an explicit means of estimating post-selection T-cell prevalence within a given MHC-class restriction.

### 3.3 Recognition of foreign and point-mutated self-peptides

RACER, which leverages structural information to assess binding strength, can be used to simulate the influence of selection on the resulting T-cell repertoire and, hence, on the recognition of TANs across patients and cancer subtypes. Applying our model to CDR_3_ *α*, *β* chains obtained from T-cell sequencing, together with possible TAN lists generated by deep sequencing of cancer populations could provide a rapid method of generating clinically actionable information for cancer specific TCRs in the form of putative TCR-TAN pairs, provided those TANs are similarly presented on the original MHC [50, 51]. While we focused our analysis on a single MHC restriction, our approach could also be applied to the crystal structure of another TCR-p-MHC pair, together with several known strong and weak binder candidates. More generally, our results also provide credence to the linear constitutive assumption which enables us to sum binding energies of individual residue pairs for quantifying TCR-peptide interactions [20, 14]. Moreover, the predictive accuracy of RACER can be further improved by including additional strong binders from crystal structures that are deposited in the database (Fig. 5B), thus providing a mechanism for additional refinement and improvements in predictive accuracy as more sequence and structural data become available.

The relative efficacy of targeting TANs remains an important question with significant clinical implications. We have shown that RACER can readily simulate full-scale thymic selection to produce an MHC-restricted T-cell repertoire. The overall agreement in post-selection behavior between this study and our previous theoretical analysis is reassuring for both approaches, in addition to the general properties of T-cell immunogenicity (Fig. 6D) and recognizing the balance between TCR diversity and similarity observed in experimental data (Fig. S11). Taken together, our findings suggest that thymic selection affords little to no recognition protection of peptides closely related to self, thus supporting the notion that T-cells undergoing central tolerance to thymic self-peptides are essentially memorizing a list of antigens to avoid. Given that a large class of TANs are generated via point mutations in self-peptide, this result also provides a quantitative argument for the efficacy of immunotherapies which target point-mutated neoantigens. When compared to experimentally observed TCR specificity, the identified antigen-specific T-cells highlights the power of RACER, when assigned a known epitope target, to identify a diverse set of antigen-specific TCRs within high-dimensional TCR primary sequence space. We expect this approach to accelerate therapeutic T-cell discovery by providing a quick and inexpensive screening tool that can then inform more costly confirmatory TCR repertoire sequencing and affinity tests. Currently, we have focused on predicting binding affinities of TCR-peptide pairs restricted to a particular MHC allele, offering a proof-of-principle for epitope identification. This procedure can in general be repeated for other MHC alleles and could be applied to a broad set of clinical scenarios by training on a relatively small number of the most common HLA Class-I alleles, which have been well-studied and have ample available crystal structure data. Toward this end, an immediate future goal will be to generalize RACER for predictions across MHC alleles and gene classes.

While important, studying TCR-p-MHC pairwise interactions on the scale of an entire T-cell repertoire is only one factor influencing adaptive immune system recognition. Signaling between other adaptive immune system elements (including helper T-cells and natural killer cells) and intracellular factors which influence antigen generation, abundance, and availability on the cell surface also affect recognition rates. Encouraged by the RACER model’s reasonable selection and recognition behavior, we propose this optimized framework as the first of its kind tool for tackling general questions regarding the interactions between the T-cell repertoire and relevant antigen landscape. Although we calculate static antigen recognition probabilities, the temporal tumor-immune interaction leads to dynamic co-evolution [24] reliant on the quality, abundance, and systems-level signaling of antigens [64]. In the setting of stem cell transplantation approaches, the availability of time series assessments of immune cell repertoires, self-peptides, and tumor antigens promises to inform optimal treatment strategies based on the donor immune system and host cancer population.

## 4 Methods

### 4.1 Details of the energy model used in our optimization

To evaluate the binding energies on the basis of a structurally motivated molecular energy model, the framework of a coarse-grained protein energy model, AWSEM force field [46], was utilized for calculating the binding energies between the T-cell receptors (TCRs) and the peptide displayed on top of a MHC molecule. AWSEM is a coarse-grained model with each residue described by the positions of its 3 atoms – C*α*, C*β* and O atoms (except for glycine, which does not have C*β* atoms) [46]. We used the C*β* atom (except for glycine, where the C*α* atom was used) of each residue to calculate inter-residue interactions. The original AWSEM energy includes both bonded and non-bonded interactions.

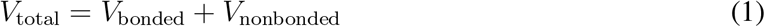

Since those residue pairs that contribute to the TCR-peptide binding energy, specifically those from the CDR loops and peptides, are in separate protein chains, only non-bonded interactions are considered. *V*_nonbonded_ is composed of three terms:

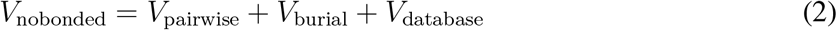

Among them, *V*_burial_ is a one-body term describing the propensity of residues to be buried in or exposed on the surface of proteins. *V*_database_ is a protein sequence-specific term that uses information from existing protein database, such as secondary and tertiary interactions, to ensure locally accurate chemistry of protein structure. Since the TCR-p-MHC system features pairwise interactions between a TCR and its corresponding peptide, only the term *V*_pairwise_ is used for this study.

The pairwise energy of AWSEM potential describes the interactions between any two non-bonded residues and can be further separated into two terms:

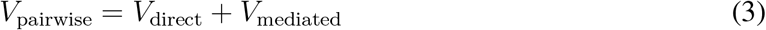

*V*_direct_ captures the direct protein-protein interaction of residues that are in between 4.5 and 6.5 Å. The functional form of *V*_direct_ is

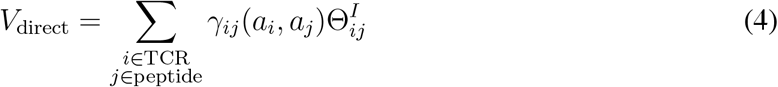

in which 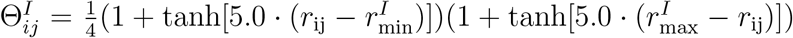 is a switching function capturing the effective range of interactions between two residues (here taken between 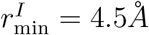 and 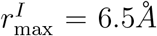). Thus, two residues are defined to be “in contact” if their distance falls between 4.5*Å* and 6.5 *Å*. *γ_ij_* (*a_i_, a_j_*) describes the residue-type dependent interaction strength, and is the most important parameter that enters the optimization of RACER. *V*_mediated_ describes the longer range interactions of two residues and is not used in this study.

### 4.2 Optimization of RACER to maximize specificity of TCR-peptide recognition

For each interaction type, the *γ_ij_*(*a_i_, a_j_*) parameters constitute a 20-by-20 matrix of parameters that describes the pairwise interaction between any two residues *i*, *j*, each with one of the 20 residue types, *a_i_*, *a_j_*. Guided by the principle of minimum frustration [43], *γ_ij_*(*a_i_, a_j_*) was previously optimized self-consistently to best separate the folded states from the misfolded states of proteins. Distilled into mathematical details, the energy model was optimized to maximize the functional *δE*/Δ*E*, where *δE* is the energy gap between folded and misfolded proteins, and Δ*E* measures the standard deviation of the energies of the misfolded states. An energy model was optimized based on a pool of selected protein structures [65], where a series of decoy structures were generated by either threading the sequences along the existing crystal structures, or by biasing the proteins into molten globule structures using MD simulations [45]. The resultant *γ* parameter thus determines an energy model that facilitates the folding of proteins with given sequences.

Motivated by this idea, RACER was parameterized to maximize the Z-scores for fully separating TCR strong binders from weak ones. Strong binders were chosen to be those top peptides that survive and were amplified to contain to at least 50 copies after four rounds of experimental deep sequencing processes (details in Section Data used in our analyses) [27], together with the peptides present in the deposited crystal structures [53]. In the experiment of [27], to ensure the correct display of peptides on the MHC, limited diversity was introduced for most distal residues and anchoring residues of peptides. The decoy binder sequences were generated by randomizing the non-anchoring residues of each strong binder thereby generating 1000 copies, and excludes the strong-binder sequences. The *γ* parameters were then optimized to maximize the stability gap between strong and randomized set of decoy binders, *δE = A*^T^*γ*, and the standard deviation of decoy energies, Δ*E*^2^ = *γ*^T^*Bγ*, where the vector A and matrix B are defined as:

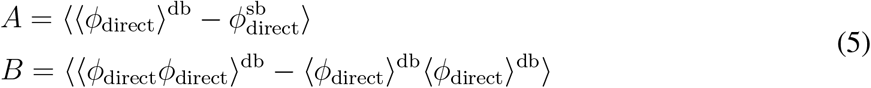

In the above Eq. 5, “direct” stands for the interaction type, *V*_direct_. *ø*_direct_ is the functional form for *V*_direct_. *ø*_direct_ also summaries the probability of contacts formation (interaction matrix) between pairs of amino acids in a specific TCR-peptide system. The subscripts “db” stands for “decoy binders” and “sb” stands for “strong binders”. The first average is over the 1000 decoy binders generated from one specific strong binder. The second average is over all the strong binders. The maximization of 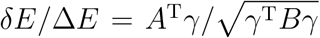 can be performed effectively by maximizing the functional objective *R*(*γ*) = *A*^T^*γ* − *λ*_1_Δ, where Δ^2^ = *γ*^T^*Bγ*. The solution of this optimization gives *γ* α *B^−1^A*. A is a vector containing the difference in the number of interactions of each type in the strong and decoy binders. B is a covariance matrix, which contains information about which types of interactions tend to co-occur in the decoy binders. Finally, *γ* is a vector that encodes the optimized strengths of the interactions. The dimension of the vector A is (1, 210), that of the matrix B is (210, 210), and that of the vector *γ* is (210, 1). To aid visual presentation, we reshape the *γ* vector into a symmetric 20 by 20 matrix in Fig. 3C. Finite sampling of decoy binders introduces noise in the optimization process, particularly in B. As such, a filter is applied to reduce the effects of the noise. The filtering scheme was performed by first diagonalizing the B matrix such that *B^−1^* = *P* Λ^*−*1^P ^−1^, where P is composed of the eigenvectors of B and Λ is made up of B’s eigenvalues. The first N modes of B (sorted in descending order by eigenvalue) are retained and the other (210 - N) eigenvalues in Λ are replaced with the Nth eigenvalue of B. The final result is robust to the choice of N. In practice, N is chosen so that no eigenvalue is close to zero. The Wolynes group performed this optimization in an iterative way where the optimized parameters were used for generating a new set of decoy protein structures [66]. In this study, since different peptides are structurally degenerate on top of MHC as observed from experiments [27], only one round of optimization was performed. Since the optimization leaves a scaling factor as a free parameter, throughout this manuscript, the binding energies are presented with reduced units. To obtain binding energies that have physical units, the scaling factor can be further calibrated to fit the experimentally determined binding affinities, such as the *K*_d_ values measured by SPR experiments (Fig. 2C).

### 4.3 Data input used in our analyses

A deep-sequencing technique was developed to assess the binding affinity of a diverse repertoire of MHC-II-presented peptides towards a certain type of TCR [27]. Specifically, 3 types of TCRs: 2B4, 5CC7 and 226, were used for selecting peptides upon four rounds of purification. The peptides that survived and enriched with multiple copies bind strongly with the corresponding TCR. In contrast, the peptides present initially but become extinct during purification represent experimentally determined weak binders. For each of the 3 TCRs, the peptides that end up with more than 50 copies after the purification process, together with the peptides presented in the crystal structures, were selected as strong binders. 1000 decoy sequences were generated for each of the strong binders by randomizing the non-anchoring residues. Both strong binders and decoys were included in the training set. In addition, to test the performance of RACER, peptides having at least 8 copies initially but disappearing during purification were selected as experimentally determined weak binders and were assigned to the test set for each TCR. To test the transferability of the model, we used weak-binding peptides of two different TCRs (e.g., 5CC7 and 226) as additional test sets distinct from the TCR used in training (e.g., 2B4).

When structural data for a specific TCR-peptide pair of interest is unavailable, we built the structure by homology modeling [67], based on a known TCR-peptide crystal structure incorporating the same TCR. Since potential steric clashes after switching peptide sequences may disfavor the strong binders used in our training set, we used Modeller [67] to refine the structures located at the TCR-peptide interface of strong binders before including them in the training process. Likewise, the binding energies of the experimentally determined weak binders were also evaluated after structural relaxation. The structural relaxation adds several seconds of computational time for each TCR-peptide pair, and thus poses a challenge for large scale repertoire analysis. However, the coarsegrained nature of RACER framework may significantly reduce the probability of side-chain clashes after switching peptide sequences. To test the accuracy of our model prediction without structural relaxation, we calculated the binding energies of strong and weak binders of TCR 2B4 by only switching the peptide sequences, omitting any structural adjustment. Our result (Fig. S12) shows comparable accuracy in separating strong from weak binders, similar to that reported in Fig. 2A. In the same vein, the transferability of RACER was also maintained without structural relaxation (Fig. S8). Encouraged by the accuracy of our coarse-grained model without relaxation, we modeled large pairwise collections of TCR-peptide interactions by only altering their corresponding sequences. For blind assessment of TCR transferability, we ask whether we can improve prediction accuracy if there are available strong binders determined in crystal structures of the target TCRs. To test this, we added interaction matrices calculated from the crystal structures of the other two TCRs as two additional strong binders in the training set. For example, in the case of TCR 2B4, the interaction matrices from the crystal structures of TCR 5CC7 and 226 were added into the training set of TCR 2B4, constituting a total of 46 strong binders. The test shows a significant improvement in predicting the binding specificity of TCR 5CC7 and 226 (Fig. 5B).

For an additional independent test of the transferability of RACER under the same MHC allele, we used the benchmark set reported in [56]. Four crystal structures are curated in their benchmark set, including three TCRs: 3QIB (2B4), 3QIU (226), 4P2Q (5CC7) and 4P2R (5CC7). Each of them have one strong-binding peptide presented in the crystal structure, and 4 weakly binding peptides. All the TCR-peptide pairs are associated with MHC-II allele IE^k^, and three of them overlap with the main dataset reported in [27]. We therefore used the energy model previously trained from TCR 2B4 to test its transferability for the other three TCR-peptide pairs. The calculated binding energies were converted into a Z score by referencing to a set of 1000 randomized peptides of corresponding TCRs: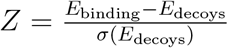, with *σ*(*E*_decoys_) being the standard deviation of *E*_decoys_. The ROC curve and AUC score were calculated by scanning through different thresholds of the Z score. A further test by including more examples from [56] is available at Supplementary note S5, Fig. S6 and S7.

### 4.4 Accuracy of RACER predictions omitting the crystal structure of target TCR-peptide pairs

To test the transferability of RACER without requiring any measured structure for a new TCR, we threaded the sequences of the CDR3 loops of the new TCR on the TCR structure used in our training. The length of CDR3*β* chain is the same among three TCRs (2B4: ASSLNWSQDTQY; 5cc7: ASSLNNANSDYT, 226: ASSLNNANSDYT), but the length of CDR3*α* chain is different (2B4: AALRATGGNNKLT; 5cc7: AAEASNTNKVV; 226: AAEPSSGQKLV). In order to accommodate such difference when threading the CDR3*α* sequences, we used a simple approach: aligning them based on the first two AA residues, leaving two gaps for TCR 5cc7 and 226. Modeller[67] was used to build the new loop structure based on these aligned new sequence, using the single structure in the training set as the template. These homology-modeled structures were then used for calculating the binding energies of the strong and weak binders of the new TCRs, using the trained interaction matrix. We also omitted the step of structural relaxation when replacing a new peptide sequence on the built structure. Such approach is unlikely to reduce RACER’s performance, as demonstrated in Fig. S12.

### 4.5 The leave-one-out cross validation

The Leave-one-out cross validation (LOOCV) was used to test the predictive power of RACER on its ability to identify strong binders. Specifically, one of the 44 strong binders of TCR 2B4 was removed from the training set, and its predicted binding energy *E*_pred_ was compared with the experimentally determined weak binders. If the median of the weak binders’ binding energies is larger than *E*_pred_ (a larger binding energy is associated with smaller affinity), the testing strong binder is successfully identified. Similar tests were performed for TCR 5cc7 and TCR 226. The performance of RACER is compared with that from the clustering of peptide sequences using the algorithm from CD-Hit [68] (See Supplementary note 1 for details).

### 4.6 Comparing the correlation of binding energies with the *K*_d_ from SPR experiments

Surface plasmon resonance (SPR) was performed to assess the binding affinities of the three TCRs towards 9 selected peptides [27]. The correlation between the predicted binding energies from RACER and the dissociation constant *K*_d_ evaluated from the SPR experiments thus constitutes a separate set of tests for the accuracy of RACER. We first built a relaxed structure with Modeller [67] for each of those TCR-peptide pairs, using the corresponding TCR structure as the template. We then used the optimized energy model of the corresponding TCR to evaluate the binding energy of each of those TCR-peptide pairs. The *K*_d_ values were obtained from fitting the SPR titration curves (Fig. S4F of [27]) using equation 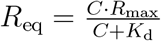 with C, *K*_d_ and *R*_max_ as free parameters. The Pearson correlation coefficient and the Spearman’s rank correlation coefficient between *k*_B_*T* log(*K*_d_) and predicted binding energies were used to quantify this correlation.

### 4.7 Evaluation of contact residues of MHC-restricted TCR-peptide pairs

The contact map of a given TCR-peptide structure was constructed by measuring the proximity W_i,j_ between each residue of peptide (residue i) and CDR loops (residue j) based on their mutual distance, using a smoothed step function:

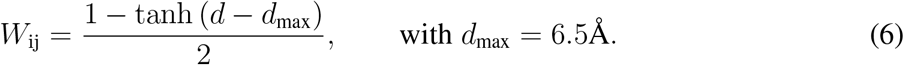

Only C*_β_* atoms were included in our calculation (except for glycine, where the C*α* atom was used). The CDR3 loops were utilized as defined in the IEDB database [69]. The constructed contact map represents those residues that are spatially close to each other in the given crystal structure.

### 4.8 Evaluation of different TCR-p-MHC interactions used for statistical study

In order to assess the statistical behavior of the inferential model, we calculated the pairwise binding interactions between a simulated T-cell population of size *N_t_* and collection of *N_n_* = 10^4^ thymic self-peptides. For this proof-of-principle study, we used TCR 2B4 as an example, uniformly varying the 10^4^ amino acids of the peptides, as well as those residues from the TCR that are in spatial contact with the peptide. TCR-peptide pairwise energies were calculated for *N_t_* = 10^5^ randomized TCR sequences using the RACER energy matrix optimized for TCR 2B4, and *N_t_* = 2000 for each of the TCR-p-IE^*k*^ systems given in Fig. 4 using energies weighted according to their contact maps, along with a model using a contact map with diagonal interactions (Fig. 6A). Substitution of TCR-peptide sequences with the newly generated ensemble yielded a total of *N_t_* * *N_n_* (10^9^ in the 2B4 case; 2 * 10^7^ for each of the cases involving the TCR-p-IE^*k*^ and diagonal contact maps) TCR-peptide pairs representing interactions occurring during thymic selection. Given our previous results (Fig. S12), we avoid the computationally expensive task of structural relaxation, and instead calculate pairwise interactions with the original structure, requiring 5,000 CPU hours on an Intel(R) Xeon(R) CPU E5-2650 v2 for the large-scale 2B4-optimized simulation.

#### 4.8.1 Thymic selection

Each T-cell survives if the maximal interaction over all self-peptides does not exceed some upper threshold. Selection thresholds were chosen to achieve 50% [11]. In all cases, the RACER-optimized energy matrix was used for energy assignment. Thymic selection was performed for each of the TCR-p-IE^*k*^ examples and their corresponding contact maps given in Fig. 4 (Fig. 6A). For each TCR-p-IE^*k*^ example, *N_t_* = 2000 pre-selection TCRs were created by varying uniformly the original TCR CDR3 *α* and *β* sequences over amino acid space, keeping the sequence lengths unchanged. A similar randomization yielded *N_n_* = 10^4^ randomized peptide sequences representing self-peptides. For each of the 2000 randomized TCRs, binding energies were calculated against the 10^4^ self-peptides by selecting the corresponding entries in the RACER-optimized energy matrix weighted by the original TCR-p-IE^*k*^ contact maps, and the maximum energy was recorded. The fraction of TCRs whose maximal binding energy exceeded the selection threshold *E_n_* traces the survival curves. This procedure, utilizing the RACER-optimized energy matrix, was repeated for a simplified model that utilizes only adjacent contacts (i.e. a strictly diagonal contact map with each entry having weight one) in the TCR-peptide interaction. The number of diagonal elements in the diagonal contact model was taken to be 20 (10 for each of the CDR3*α*-peptide and CDR3*β*-peptide pairs).

#### 4.8.2 Self-peptide potency

Most self-peptides present in thymic selection are expected to participate in the deletion of self-reactive T-cells. Thus, a reasonable model of thymic selection would feature a majority of self-peptides contributing to the selection of immature T-cells. A rank order of these self-peptides based on their ability to recognize unique T-cells, or potency, characterizes the extent to which each self-peptide is utilized in thymic selection. The rank order of potency was created for the RACER model utilizing the crystal structure of the 2B4 TCR (PDB ID: 3QIB) and its corresponding energy matrix derived from the set of experimentally determined good-binders. The thymic selection process using 10^4^ self-peptides and 10^5^ TCRs for the 2B4-optimized RACER model described above generates a total of 10^9^ pairwise binding energies. The negative selection threshold *E_n_* was selected to yield 50% selection, resulting in ~ 5 . 10^4^ deleted TCRs. The number of TCRs deleted by each self-peptide was recorded. The peptide deleting the most TCRs defines the most potent self-peptide. TCRs recognized by this peptide are removed from the list of total TCRs, and this peptide is similarly removed from the list of self-peptides. This process is repeated on the smaller TCR and self-peptide list to determine the second most potent peptide. Additional iteration until no TCRs remain provides the rank order of self-peptides in decreasing order of potency. The cumulative fraction of deleted relative to total TCRs is plotted in decreasing order of peptide potency.

#### 4.8.3 Antigen recognition probabilities for individual T-cells and T-cell repertoires

Utilizing the same post-selection T-cell repertoire from the previous section, post-selection T-cells were quantified for their ability to recognize random non-self-antigens and tumor neoantigens that differ from one of the *N_n_* thymic self peptides by one residue. 50% selection of TCRs result in approximately 5 . 10^4^ surviving, for which pairwise interactions are generated against 10^3^ random and 10^3^ point-mutated self-peptides, representing foreign and tumor-associated neoantigens, respectively (randomly generated peptides were checked to ensure non-membership in the set of thymic self-peptides). Estimates of individual TCR recognition probability were calculated by averaging the 5 . 10^4^-by-10^3^ indicator matrix, having values of 1 (resp. 0) corresponding to recognition (resp. no recognition). The previous quantity estimates an individual TCR’s antigen recognition ability. Estimates of the corresponding recognition probability for the entire post-selection MHC-restricted T-cell repertoire was calculated by assessing the 1-by-10^3^ vector indicating the presence or absence of at least 1 recognizing TCR. The post-selection individual and repertoire T-cell recognition probabilities of random and point-mutant antigens were then compared with previously derived analytic results for two random energy models [20].

## 5 Acknowledgments

Work at the Center for Theoretical Biological Physics was sponsored by the NSF (Grant PHY-2019745). HL was also support by the NSF (Grant PHY-1935762). JNO was also supported by the NSF (Grant CHE-1614101) and the Welch Foundation (Grant C-1792). JTG was supported by the National Cancer Institute of NIH (F30CA213878). JNO is a CPRIT Scholar in Cancer Research.

## 6 Data Availability

The data comprised of the peptides recognized by the three TCRs, used for RACER training and testing, are available from [27]. An extended data set of these three TCRs were uploaded at Github: https://github.com/XingchengLin/RACER.git. The additional data used for training and testing on different MHC-II TCRs can be found in [56]. All other output from this study are available from the corresponding author upon reasonable request.

## 7 Code Availability

The full code, along with a demo for predicting TCR-peptide interaction, as well as being applied to a collection of randomly-generated TCRs and peptides, can be found at https://github.com/XingchengLin/RACER.git

## Footnotes

1 The Z-score is defined as the difference between the average binding energies of strong binders versus decoys, divided by the standard deviation of the decoy energies. Throughout this manuscript, we report the absolute value of the calculated Z-score, except for Fig. 5C.

## Notes

### Competing Interest Statement

The authors have declared no competing interest.

### Summary of Updates

Updated manuscript presentation; cross-validation on independent test datasets; explicit mappings to physical T-cell-peptide contact maps.

