## Supplementary Information for "Rapid Assessment of T-Cell Receptor Specificity of the Immune Repertoire"

### Supporting Information for Rapid Assessment of T-Cell Receptor Specificity of the Immune Repertoire

[1]Center for Theoretical Biological Physics, Rice University, Houston, TX
[2]Department of Physics and Astronomy, Rice University, Houston, TX
[3]Department of Chemistry, Massachusetts Institute of Technology, Cambridge, MA
[4]Medical Scientist Training Program, Baylor College of Medicine, Houston, TX
[5]Departments of Chemistry, Rice University, Houston, TX
[6]Department of Physics, Northeastern University, Boston, MA
[7]Koch Institute for Integrative Cancer Research, Cambridge, MA
[8]Department of Biological Engineering, Massachusetts Institute of Technology, Cambridge, MA
[9]Ragon Institute of MIT, MGH, and Harvard, Cambridge, MA
[10]Department of Physics, Freie Universität, Berlin, Germany
[11]Department of Biosciences, Rice University, Houston, TX
[*]Equal contribution

#### Contents

**S1 Sequence-based peptide clustering** **2**

**S2 Additional hold-out tests on an extended dataset** **2**

**S3 Sequence diversity in the leave-one-out test** **2**

**S4 The standard protein force field cannot fully resolve strong binders from weak binders** **2**

**S5 Extended test of RACER's transferability across different TCRs restricted to the same MHC-II allele** **3**

**S6 Comparison with ERGO [1], a sequence-based predictor trained by neutral network** **3**

**S7 Supplementary Figures S1-S12 and Table S1** **3**

### S1 Sequence-based peptide clustering

To assess the ability to separate strong binders from weak binders based only on the peptide sequences, CD-Hit [2, 3], a greedy incremental clustering algorithm, was used for classifying the peptide sequences of TCR 2B4. The input sequences contained a total of 44 experimentally determined strong binders (including the native peptide present in the crystal structure), as well as 231 experimentally determined weak binders as described in the main text. A strong binder was identified if it is classified in the same category of the native sequence. The best performance of CD-Hit correctly identifies the cluster including 19 strong binders (native peptide included) and no weak binders, using a sequence identity threshold of 0.5.

### S2 Additional hold-out tests on an extended dataset

To test the limit of RACER's transferability over a more diversified coverage of peptide sequence, we included more strong binders from the [4], where all peptides of TCR 2B4 that ends up with more than one copy from the deep-sequencing experiments were included, constituting an extended dataset. RACER was applied on a more demanding set of hold-out tests on the original dataset used in the main text, as well as this more extended dataset.

In the leave-50%-out-test, these strong binders were randomly shuffled before partitioned into two sets. One set was used in the training set, the other set, together with the experimentally determined weak binders, were used as the testing set. We switched two sets of strong binders for an equivalent testing, therefore constituting two testing cases for each data set. As shown in Fig. S2, the leave-50%-out-test demonstrates that RACER can fully separate strong binders from the weak ones, with an average recognition Z-score equal to 5.26 for the original dataset, and 4.63 for the extended dataset.

In the leave-90%-out-test, the strong binders were randomly shuffled into 10 sets, and we only used one of them for training, the other 9 sets, together with the experimentally determined weak binders, were used as the testing set. Therefore, we have 10 testing cases for each dataset. As shown in Fig. S2, the leave-90%-out-test again demonstrate RACER's success in distinguishing strong binders from weak ones in both the original dataset (average Z-score of 4.74) and the extended dataset (average Z-score of 4.50).

To push the limit of RACER's predictive power, we added an additional leave-99%-out test for the extended dataset, where $\sim 4$ strong binders were included in the training set, and a leave-one-in test, where only one strong binder was included in the training set. We summarize the percentage of strong peptides that failed to be detected (failure is defined to occur whenever the binding energies of the withheld binders are larger than the median of the weak binders). As shown in Fig. S3, the amount of peptides that can be recognized by RACER gradually decreases as fewer peptides were included in training, and if we only include one peptide in our training, the performance of RACER is worse than an alternative test that uses the identity of sequences based on the native peptide of the crystal structure (PDB ID: 3QIB).

### S3 Sequence diversity in the leave-one-out test

To see the coverage of sequence diversity of peptides that succeeded or failed to be recognized by RACER, we calculated the sequence identity of the peptide sequences that were used in our leave-one-out test of the TCR 2B4, based on the native peptide presented in the crystal structure (PDB ID: 3QIB). As shown in Fig. S1, RACER capably recognizes strong binders with small sequence identity in both the original dataset and the extended dataset, with some cases having little to no sequence identity.

### S4 The standard protein force field cannot fully resolve strong binders from weak binders

Two commonly used force fields were utilized to test the performance of standard protein force fields for distinguishing strong TCR binders from weak ones. This analysis was applied to the three TCRs investigated in the main text. The default AWSEM force field [5] was previously optimized for folding protein structures [6]. The Miyazawa-Jernigan (MJ) potential [7] is one of the most widely used knowledge-based force fields, derived from a statistical analysis of large protein repositories. Both force fields have been demonstrated to perform well in describing the structural dynamics of generic proteins. However, when replacing our optimized TCR parameters with these two force fields, the resulting model is unable to clearly separate the strong binders from the weak ones. (Fig. S4).

### S5 Extended test of RACER's transferability across different TCRs restricted to the same MHC-II allele

To test the transferability of RACER beyond the coverage of the TCR-p-MHCs used in the main text, we further tested the performance of RACER on the data provided in [8]. The data includes all the TCRs associated with different MHC-II alleles until 2017. One strongly binding peptide, and four weakly binding peptides were provided for each TCR. We used RACER to perform a leave-one-out prediction for all the cases where more than one TCRs are shared among the same allele, by excluding one TCR from training and using the optimized interaction matrix to predict the binding affinity of the withheld TCR. The first 50 eigenvectors of the B matrix in Eq. (5) (see Method for details) were found to be well-determined. The influence of the remaining eigenvectors of the B matrix on the optimized interaction parameters in $\gamma$ was reduced according to a filtering scheme (see Method for details). As shown in Figure S6, RACER was able to recognize the strong peptide (Z-score $> 1$) for 21 out of the 26 tests. It is worth noting that there are many cases where TCRs associated with the same MHC allele share different V-alpha and V-beta genes. RACER works less well for cases where only 2 TCRs are available, and better when there are $3 \sim 5$ cases, regardless of whether they share the same or different V-alpha and V-beta genes. This additional test further supports the predictive power of RACER trained with a small set (around 3 to 5 copies) of available TCR-peptide structures/sequences. To challenge RACER's predictive capacity when statistical learning is performed on a TCR-pMHC pair distinct from either the target predicted TCR or peptide, we intentionally selected a TCR-pMHC structure with the same MHC allele having different V-alpha and V-beta genes, where available, from the target as the template. We replaced the CDR3 loops with target sequences using trivial alignment (case III of Fig. 1), and repeat the same test as above. As shown by Fig. S7, RACER can still recognize 19 out of the 26 examples (Z-score $>$ 1). The success of this test highlight RACER's predictive power even if only one crystal structure of the same allele is utilized.

### S6 Comparison with ERGO [1], a sequence-based predictor trained by neutral network

ERGO [1] is a sequence-based TCR-peptide prediction tool trained by neutral networks. ERGO implemented two types of models: Long short-term memory (LSTM) and Autoencoder, together with two training datasets: McPAS-TCR ($2 \times 10^4$ TCR by 300 peptides) and VDJdb ($4 \times 10^4$ TCR by 200 peptides). We applied ERGO to calculate the binding scores of the strong and weak binders of the three TCRs in [4]. We found ERGO performs best when the Autoencoder was applied based on the VDJdb database. As shown in Fig. S8, ERGO can only recognize the strong binders of TCR 5CC7 with a Z-score of 2.85.

### S7 Supplementary Figures S1-S12 and Table S1

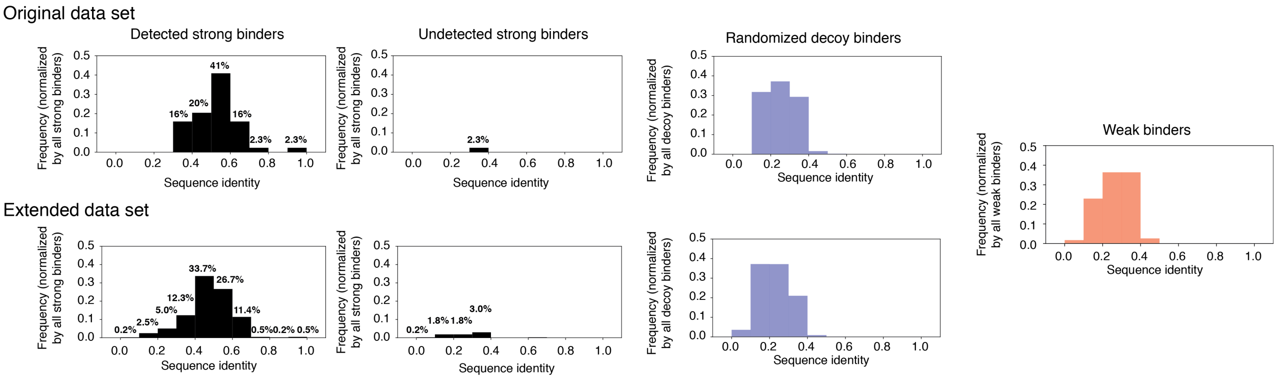


Figure S1: The normalized count frequency of the sequence identity, calculated based on the native peptide of TCR 2B4, between the experimentally determined strong binders (black), randomized decoy binders (blue), and experimentally determined weak binders (orange). For the strong binders, the probability distributions are further organized based on whether or not they are successfully detected in the leave-one-out test. For the strong binders, the normalization was carried out with the total number of strong binders being the normalization factor, to emphasize the contrast between the number of detected/undetected cases. Detailed percentage of histograms within each bin are noted. We show the distribution of sequence identities for the original dataset (including all strong binders where the final copies of peptides from the experiment were amplified by more than 50 times), and an extended dataset (including additional peptides where the final copy numbers were larger than 1).

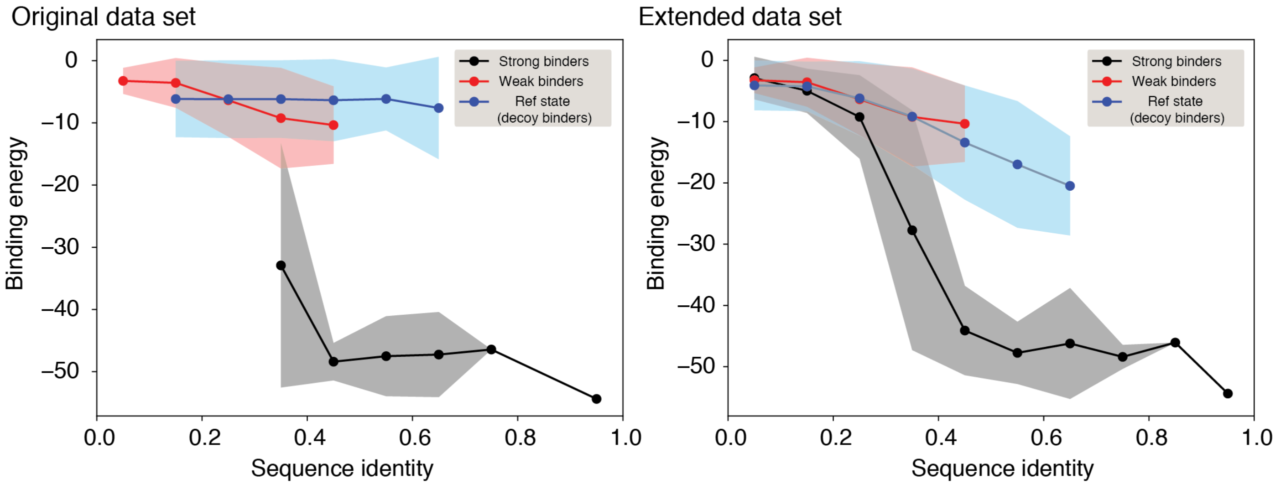


Figure S2: Cross-validation test of TCR 2B4 with RACER (main text Fig. 2) where 50% (A, C) and 10% (B, D) of the strong binders were used as the training set (blue). The predicted binding energies of the 50% of withheld strong binders (yellow) are lower than the binding energies of the experimentally determined weak binders (brown). The median of each set of binders was shown as a bar in the corresponding box plot. The whiskers are placed at the first and last datum points that fall within (m, M), where m = Q1 - 1.5IQR and M = Q3 + 1.5IQR, with IQR = Q3 - Q1 representing the interquartile range. The calculated Z-score of each test was shown at the top. In both the original (A, C) and extended (B, D) dataset, the leave-50%-out and leave-90%-out test demonstrate RACER's predictive capacity for recognizing strong binders of the TCR 2B4.

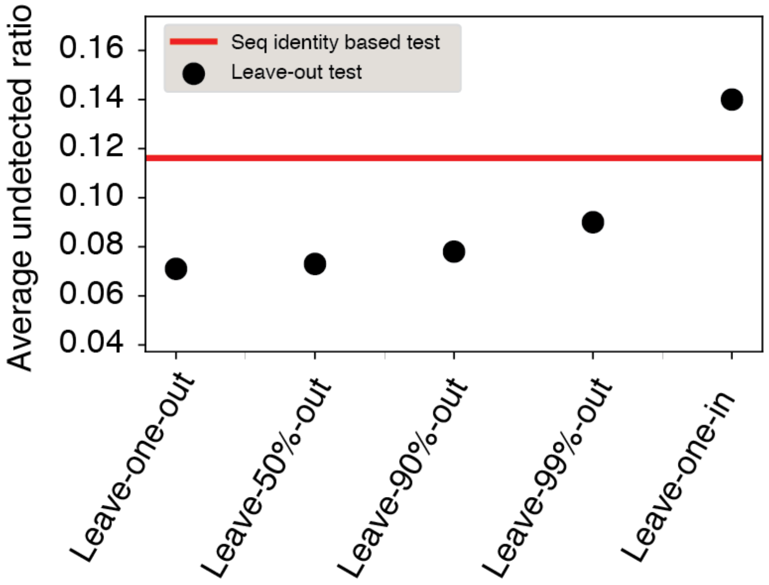


Figure S3: Cross-validation test of TCR 2B4 with RACER for the extended dataset, where more diversified strong binders were included for a more comprehensive test. RACER performs well, reliably detecting > 99% of the strong binders, until leaving 99% of the peptides out. At this point RACER's performance deteriorates due to a lack of training data. When only one peptide is included in the training set, RACER performs worse than a selection based only on the sequence identity of peptides calculated based on the native peptide in the crystal structure (PDB IDid: 3QIB) of the TCR 2B4.

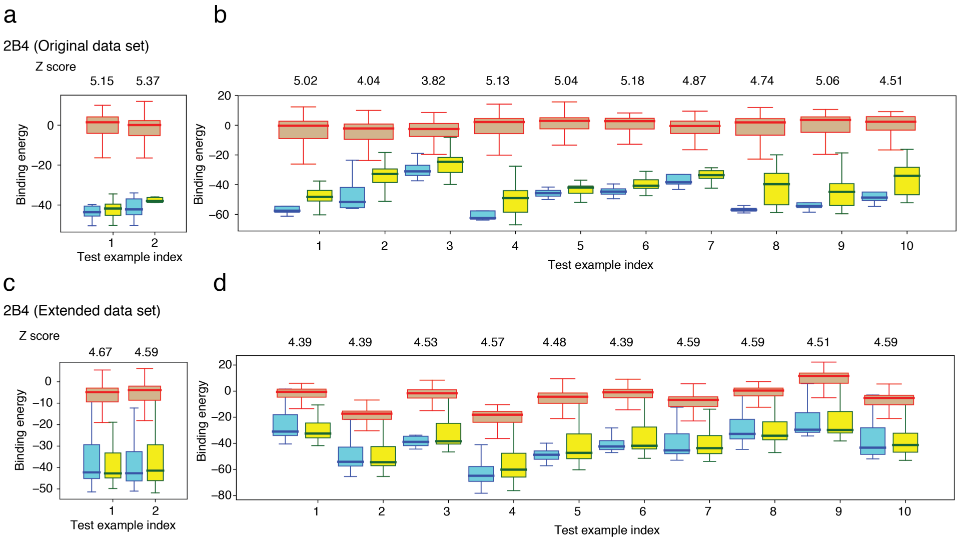


Figure S4: The protein interaction parameters from standard force field cannot fully separate strong binders from weak ones. **A,** The performance using the default parameters from the AWSEM force field [5]. **B,** The performance using the parameters from the Miyazawa and Jernigan (MJ) potential [7]. Compare this with the main text Figure 2 shows the advantage of RACER in terms of identifying strong binders from weak binders.

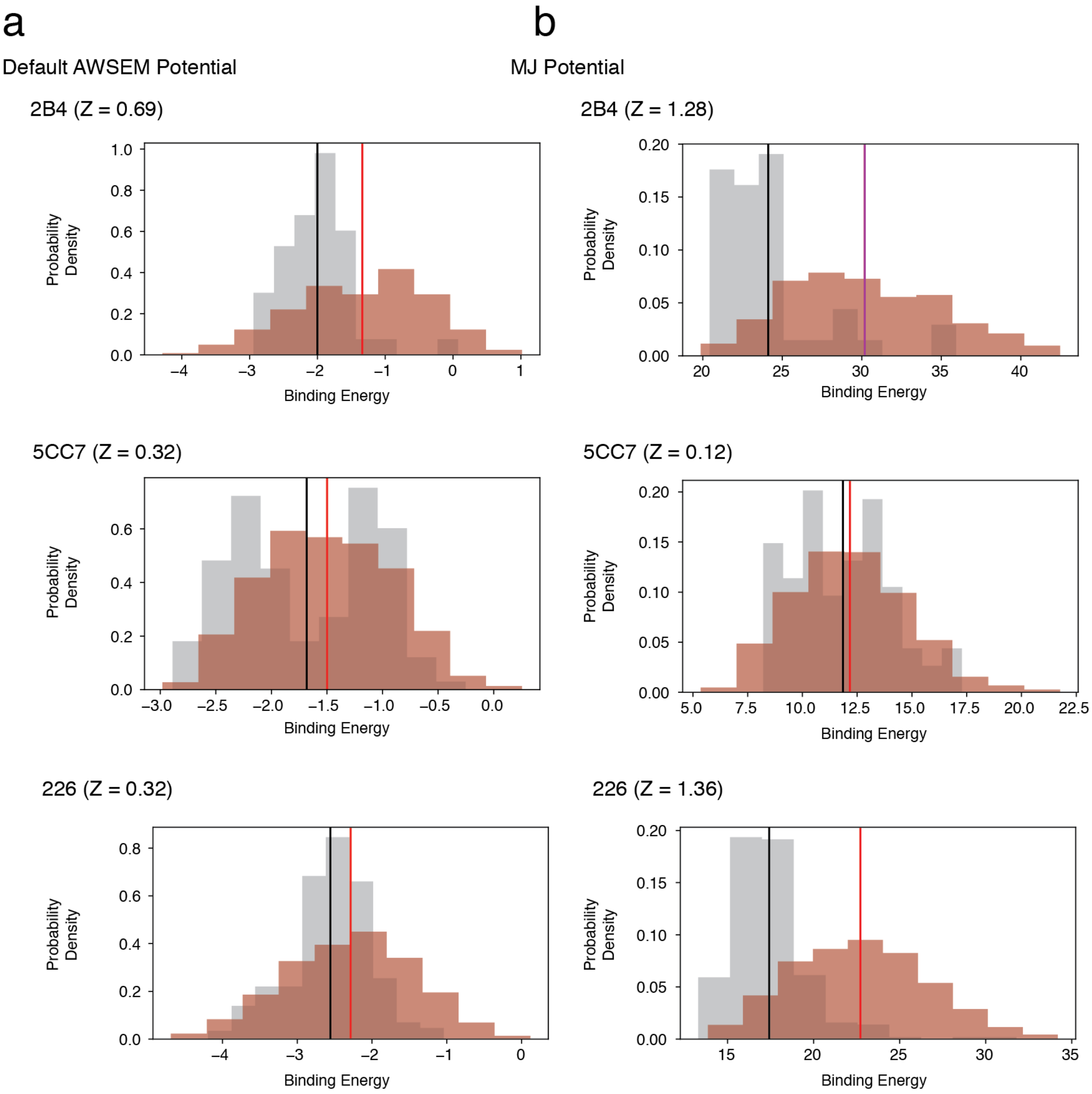


Figure S5: The contact maps of TCR-peptide pairs associated with different MHCII alleles share less structural similarity, compared with the main text Figure 4. Contact maps are calculated using distances from each pairwise TCR-peptide amino acid combination using Eq. 6 for the following TCR-p-MHC pairs: 4P2Q - peptide ADGLAYFRSSFK presented by MHC-II IE$^k$ to TCR 5cc7 **A,** CDR3$\alpha$ (AAEASNTNKVV) and **B,** CDR3$\beta$ (ASSLNNANSDYT); 3MBE - peptide AMKRHGLDNYRG presented by MHC-II IA$^g$ to TCR 21.3 **C,** CDR3$\alpha$ (AAEDGGSGNKLI) and **D,** CDR3$\beta$ (ASSWDRAGNTLY); 3C5Z - peptide FEAWKAKANKA presented by MHC-II IA$^b$ to TCR B3K506 **E,** CDR3$\alpha$ (ALVISNTNKVV) and **F,** CDR3$\beta$ (ASIDSSGNTLY).

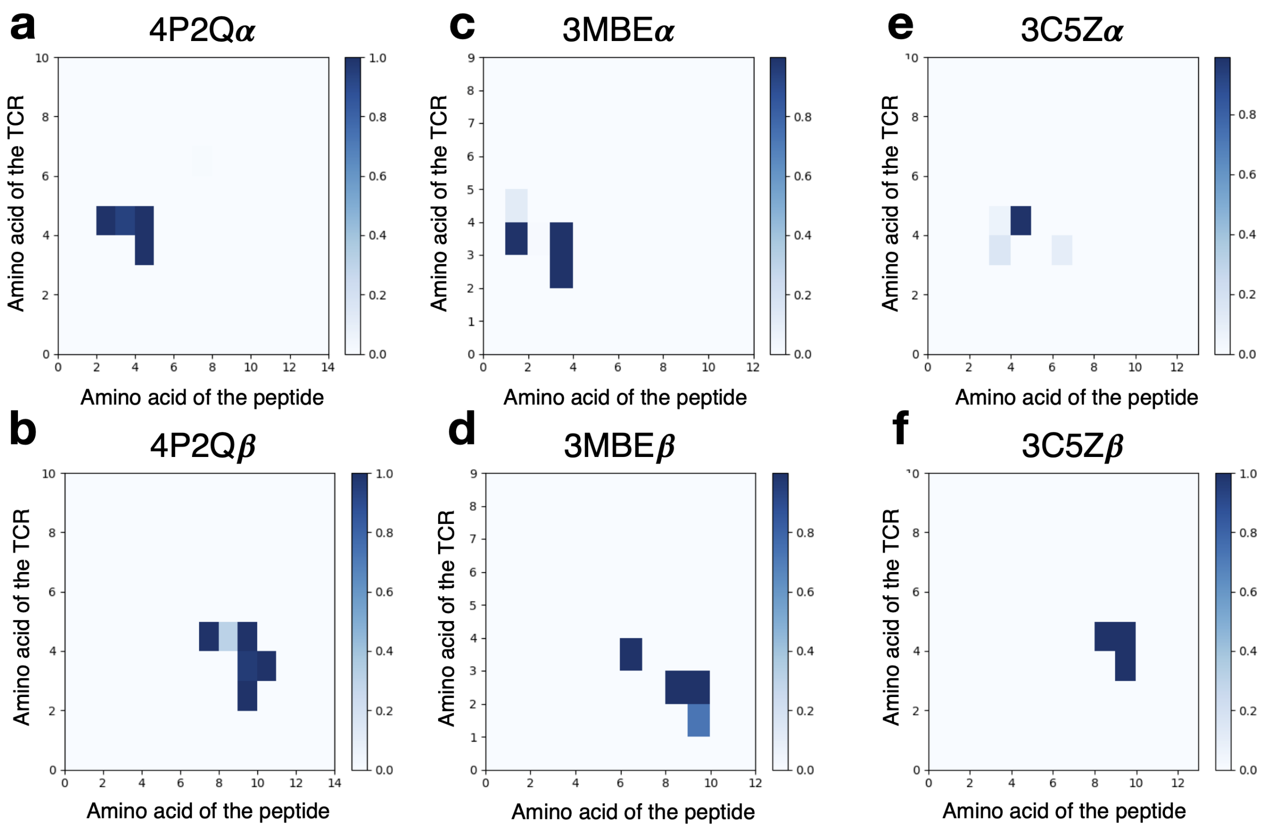


Figure S6: Further leave-one-out test of RACER's predictive transferability over TCRs, using data from [8]. The TCRs were grouped by their associated MHC allele, with their V-alpha and V-beta genes noted at the bottom. Asterisks are marked for TCRs sharing identical peptides within the same allele. RACER successfully predicted lower binding energies for strong binders (blue) relative to weak binders (brown). The prediction Z-score is provided above each case. RACER was able to successfully recognize the strong-binding peptide (Z-score $> 1$) for 21 out of the 26 tests. RACER's predictive accuracy is reduced in cases where only 2 TCRs are available, and shows improvement when there are 3∼5 cases, regardless of whether they share the same V-alpha and V-beta genes.

| TCR | Gene usage (alpha) | Gene usage (beta) | CDR sequences (alpha) | CDR sequences (beta) | Native peptide sequence |
|---|---|---|---|---|---|
| 2B4 | V:TRAV4N-4*01<br>J:TRAJ56*01 | V:TRBV26*01<br>J:TRBJ2-5*01 | CDR1:TTMRA<br>CDR2:LASGT<br>CDR3:AALRATGGNNKLT | CDR1:KGHPV<br>CDR2:FQNQEV<br>CDR3:ASSLNWSQDTQY | ADLIAYLKQATKG |
| 5CC7 | V:TRAV4N-4*01<br>J:TRAJ34*01 | V:TRBV26*01<br>J:TRBJ1-2*01 | CDR1:TTMRA<br>CDR2:LASGT<br>CDR3:AAEASNTNKVV | CDR1:KGHPV<br>CDR2:FQNQEV<br>CDR3:ASSLNNANSDYT | ANGVAFFLTPFKA |
| 226 | V:TRAV4N-4*01<br>J:TRAJ16*01 | V:TRBV26*01<br>J:TRBJ1-2*01 | CDR1:TTMRA<br>CDR2:LASGT<br>CDR3:AAEPSSGQKLV | CDR1:KGHPV<br>CDR2:FQNQEV<br>CDR3:ASSLNNANSDYT | ADLIAYLKQATKG |

Table S1: Detailed information[11] of TCR 2B4, 5CC7 and 226 used in the main text. The three TCRs used in our test shared the same V gene, their J genes are different from each other, resulting in different CDR3 sequences.

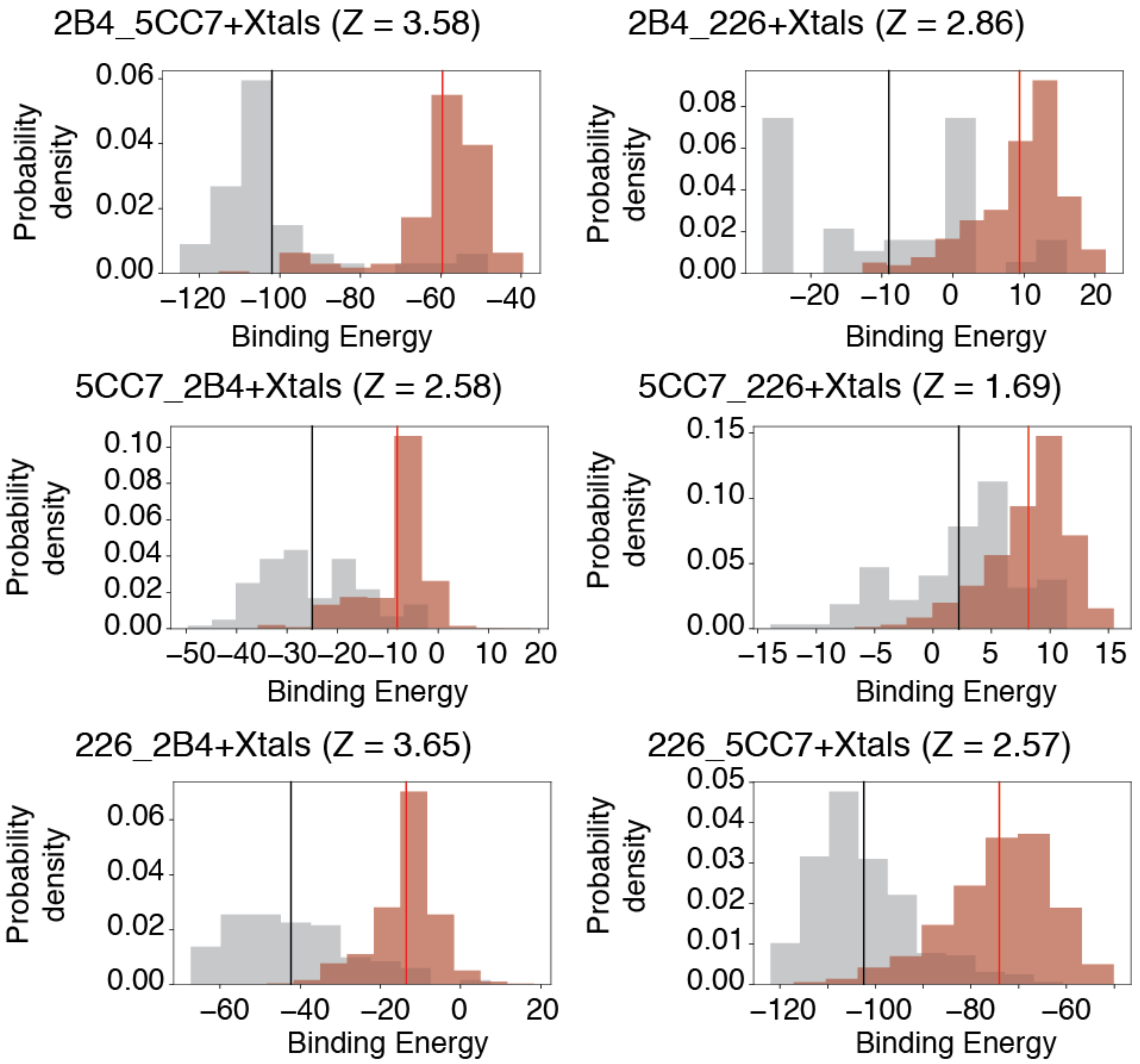


Figure S7: Additional leave-one-out test of RACER's predictive transferability over TCRs, using data from [8]. One template structure, intentionally selected to have V-alpha and V-beta genes that are distinct from the target structure (where available), is used to build the target structure; this is accomplished by replacing the CDR3 loops with target sequences based on trivial alignment (case III of Fig. 1). Asterisks mark examples where the template structure shares the same peptide sequence as the target structure. For all remaining cases target amino acid sequences are explicitly compared to the template sequence, with red (resp. cyan) positions having different (resp. identical) amino acid entries. Double-ended arrows indicate those examples where the template and the target have also been switched for another prediction test. RACER successfully predicted lower binding energies of the strong binders (blue) relative to the weak binders (brown). The TCRs were grouped by their associated MHC allele, with their V-alpha and V-beta genes, as well as the template structures labeled at the bottom. RACER-predicted Z-scores are listed at the top of each case. RACER was able to recognize the strong peptide (Z-score $>$ 1) for 19 out of the 26 challenging tests, and maintains predictive accuracy when only restricted to test cases which include distinct peptides from the training step (accuracy 67%
.

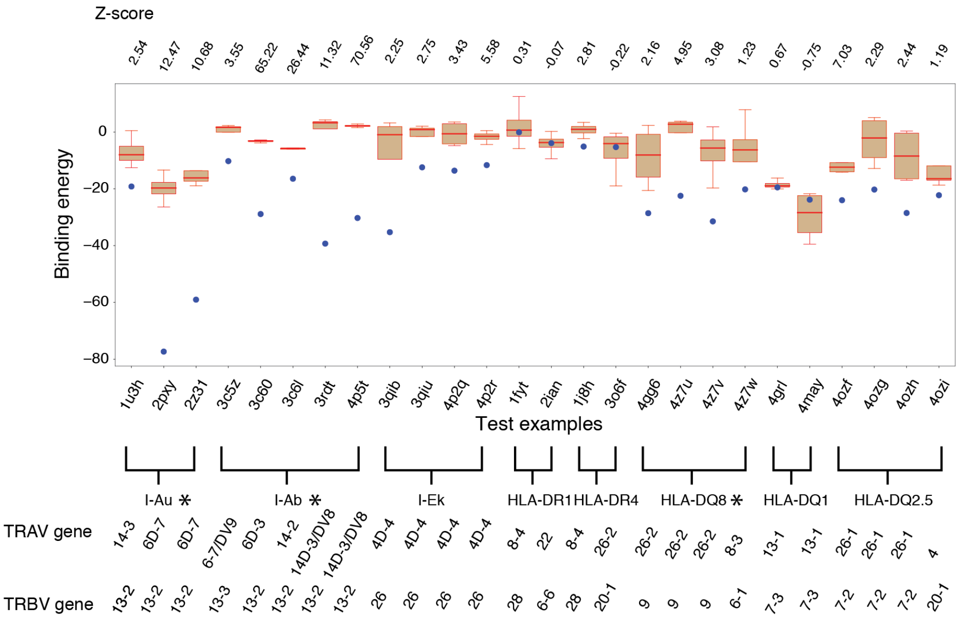


Figure S8: RACER shows transferability in terms of predicting TCR-p-MHC interactions without requiring any new structure for the new TCR. The figure is formatted in the same way as Figure 5A of main text. The energy model trained based on one TCR (e.g. 2B4) is capable of resolving the experimentally determined strong binders from weak binders of the other two TCRs (e.g., 5CC7 and 226).

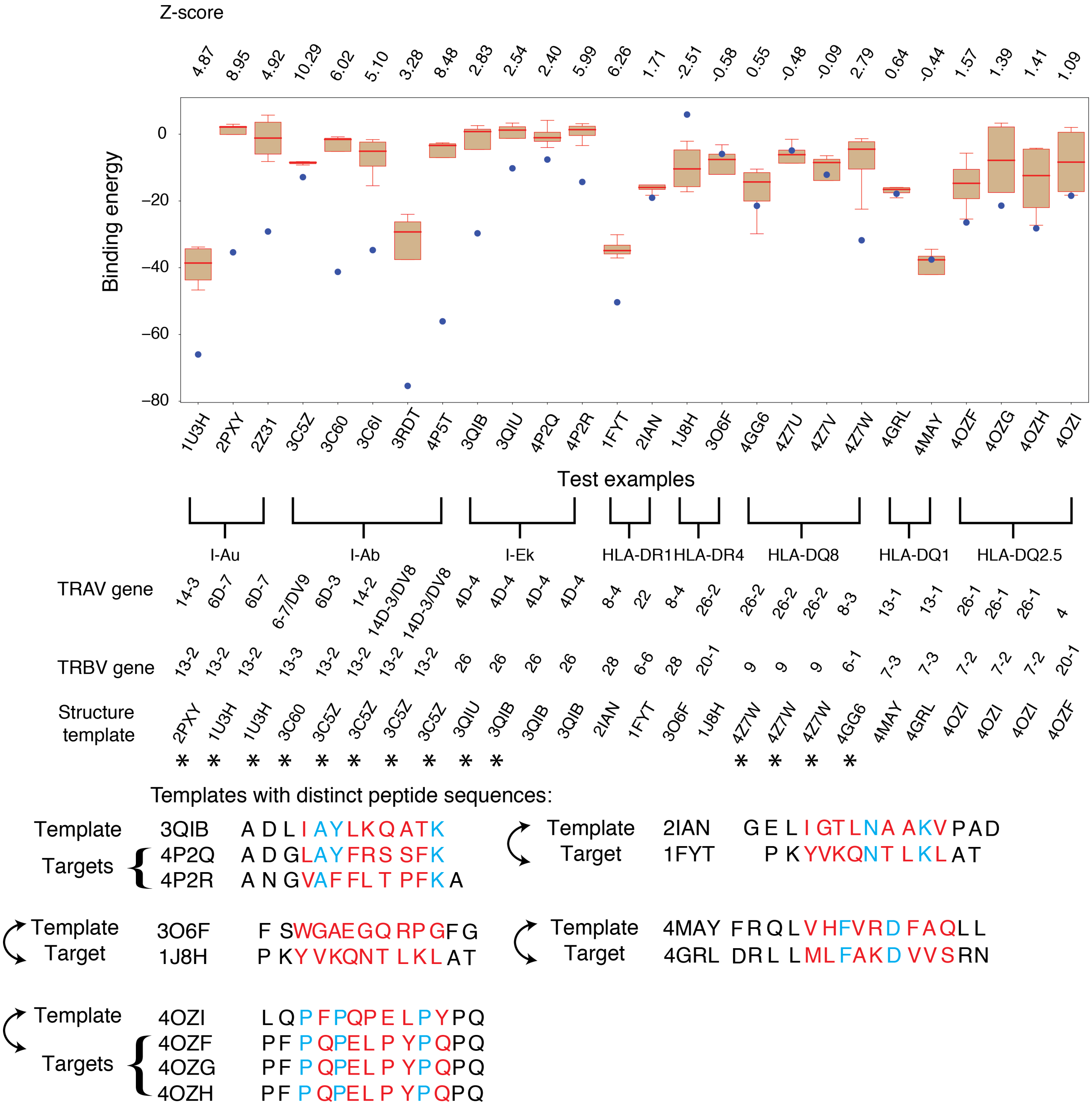


Figure S9: The performance of ERGO [1] for differentiation of the strong and weak binders of three TCRs. The best performance of ERGO, using Autoencoder trained on the database VDJdb, can recognize the strong binders of TCR 5CC7, with a recognition Z score of 2.85. However, ERGO cannot fully differentiate the strong binders of TCR 2B4 and 226.

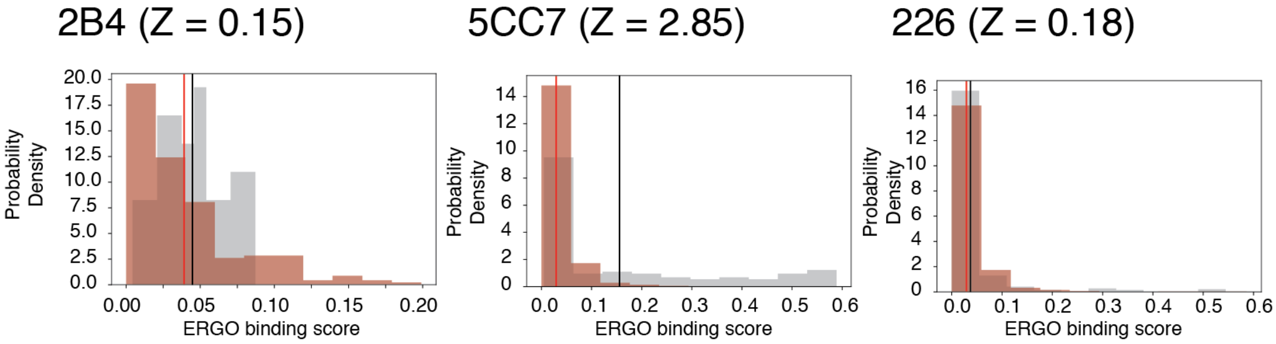


Figure S10: Comparison of RACER recognition characteristics to previous models [9]. RACER-derived estimates of post-thymic selection T-cell repertoire recognition rates reveal similarity in the ability to recognize foreign and point-mutated self antigen, in agreement with previous models. **A,** Participation of self-peptides in the deletion of reactive TCRs is quantified by plotting the total number of unique TCRs recognized as a function of each self-peptide rank-ordered based on selection potency. **B,** The probability of post-selection individual TCR recognition of foreign and point-mutated self-peptide as a function of the percentage of surviving TCRs following negative selection; positive standard deviations are given for estimates obtained in the RACER model (in all plots purple represents outputs of the RACER model, red and blue correspond to PIRA and RICE models from [9], respectively) **C,** The product of thymic selection survival probability and recognition probability of random and point-mutated self-peptides as a function of T-cells survival probability in the RACER model.

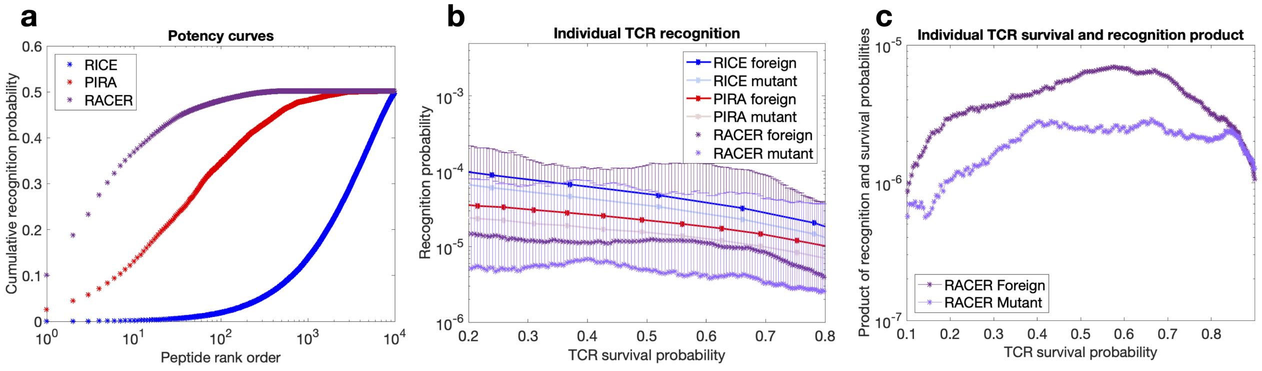


Figure S11: Quantification of the foreign epitope-specific T-cell repertoire. Each of the $10^4$ T-cells were filtered based on their ability to recognize foreign epitopes in the RACER simulation, performed for the pre-thymic selection and post-selection T-cell repertoire. Each foreign epitope was then sorted by the number of total post-selection T-cells that recognized that antigen. (A) Dendrograms of the identified post-selection recognizing T-cells for the top 10 recognized peptides are constructed using hierarchical clustering using an averaged hamming distance on the primary CDR3 sequence of each T-cell in a similar manner as in [10]. (B) For each peptide, two dissimilar TCRs were selected randomly from the left and right side of the highest dendrogram clade. Each of these TCR underwent mutagenesis by point-mutating a single entry in the CDR3 region 50 times for a total of 100 mutated (closely-related) TCRs (blue and red clusters). These TCRs were then subject to the same thymic selection and foreign peptide recognition steps as previously, and dendrograms of these and the original TCRs were constructed. (The following peptide sequences were identified by RACER as the most immunogenic: 865=ADWINQGSDWWKG, 574=ADLIALLLMWWKG, 364=ADAIAEAANCSKG, 49=ADEINKHEKWWKG, 647=ADMIDSKSTSAKG, 520=ADSIAHCGKFSKG, 386=ADWITHNWALWKG, 394=ADCIAYPKRDAKG, 550=ADYINACKSDAKG, 588=ADYINPTWAHAKG).

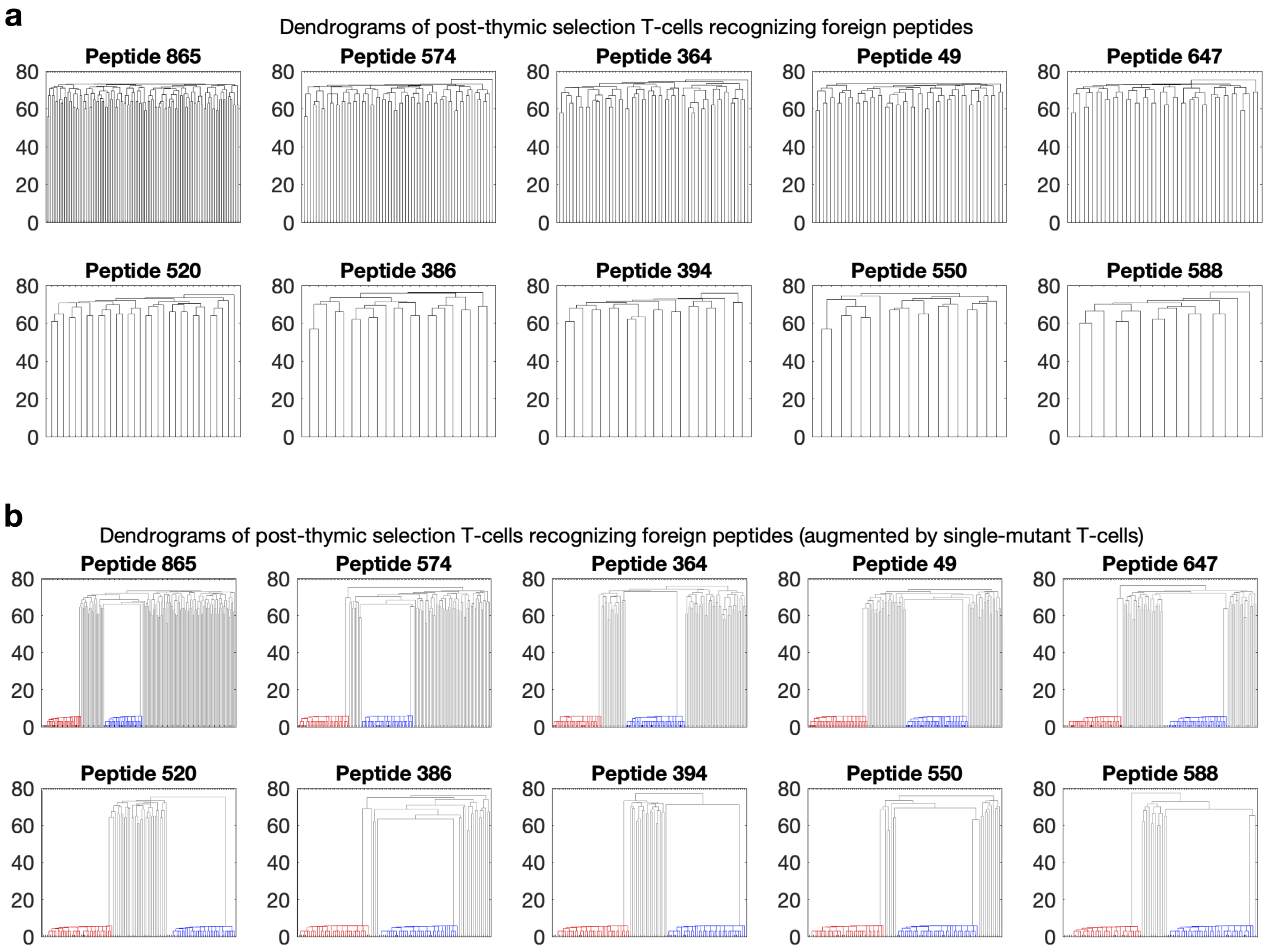


Figure S12: RACER ensures comparable accuracy with or without structural relaxation after changing peptide sequences. The binding energies of experimentally determined strong and weak binders as predicted by RACER with **A,** and without **B,** structural relaxation after switching the peptide sequences. The coarse-grained nature of RACER significantly reduces the chance for steric clashes to occur after changing peptide residues, resulting in comparable modeling performance.
